# Structural characterization and inhibition of the interaction between ch-TOG and TACC3

**DOI:** 10.1101/2024.05.31.596836

**Authors:** James Shelford, Selena G. Burgess, Elena Rostkova, Mark W. Richards, Christian Tiede, Alistair J. Fielding, Tina Daviter, Darren C. Tomlinson, Antonio N. Calabrese, Mark Pfuhl, Richard Bayliss, Stephen J. Royle

## Abstract

The mitotic spindle is a bipolar array of microtubules, radiating from the poles which each contain a centrosome, embedded in pericentriolar material. Two proteins, ch-TOG and TACC3, have multiple functions at the mitotic spindle due to operating alone, together or in complex with other proteins. To distinguish these activities, we need new molecular tools to dissect their function. Here, we present the structure of the **α**-helical bundle domain of ch-TOG that mediates its interaction with TACC3 and a structural model describing the interaction, supported by biophysical and biochemical data. We have isolated Affimer tools to precisely target the ch-TOG-binding site on TACC3 in live cells, which displace ch-TOG without affecting the spindle localization of other protein complex components. Inhibition of the TACC3–ch-TOG interaction led unexpectedly to fragmentation of the pericentriolar material in metaphase cells following the formation of a bipolar spindle and delayed mitotic progression; uncovering a novel role of TACC3–ch-TOG in maintaining pericentriolar material integrity during mitosis to ensure timely cell division.

## Introduction

Chromosome segregation during mitosis is driven by the mitotic spindle (McIntosh, 2016). To form a spindle, the centrosomes separate and move to opposite ends of the cell and a bipolar microtubule array is generated between them. The microtubules contact the kinetochores of each paired sister chromatid to coordinate their movement, while at the other end, the microtubules are focused at the spindle pole which contains a centrosome: a centriole pair embedded in pericentriolar material (Prosser and Pelletier, 2017). Understanding how the mitotic spindle assembles and operates are major goals in cell biology.

Many proteins contribute to the formation and function of the mitotic spindle; this paper focusses on two: ch-TOG and TACC3. These proteins interact with each other and participate in a number of activities (Saatci and Sahin, 2023). First, a non-motor protein complex that binds spindle microtubules (MTs) is composed of TACC3, ch-TOG, clathrin and GTSE1 (Fu et al., 2010; Hubner et al., 2010; Lin et al., 2010; Booth et al., 2011; Bendre et al., 2016). This complex stabilizes the bundles of spindle microtubules that attach to kinetochores by physically crosslinking them (Hepler et al., 1970; Booth et al., 2011; Nixon et al., 2015, 2017). The complex is formed by clathrin and TACC3 at its core, which bind to the microtubule lattice, while ch-TOG and GTSE1 bind respectively to TACC3 and clathrin as ancillary subunits (Hood et al., 2013; Burgess et al., 2018; Ryan et al., 2021). Second, ch-TOG can bind to kinetochores, independently of microtubules, via an interaction with Hec1 where it plays a role in mitotic error correction (Herman et al., 2020). Third, TACC3 and ch-TOG in an exclusive complex, track the growing ends of microtubules in the spindle (van der Vaart et al., 2012; Nwagbara et al., 2014; Gutiérrez-Caballero et al., 2015). Fourth, at the spindle pole, ch-TOG localizes to the centrosome whereas TACC3 is observed close by (Gutiérrez-Caballero et al., 2015). Understanding the relative contributions of these proteins to spindle assembly and function is therefore complicated and is hampered by a lack of molecular tools to deconvolute the roles of individual proteins cleanly.

Mammalian ch-TOG is a member of the XMAP215 family of microtubule polymerases. These proteins vary in size, with an N-terminal region comprising 2, 3 or 5 TOG domains and a C-terminal region that is variable. A TOG domain is a module consisting of 6 HEAT repeats which can bind to tubulin dimers (Ayaz et al., 2012; Fox et al., 2014). The current model for XMAP215 proteins is that they track the growing end of the microtubule and contribute to its polymerization using the multiple TOG domains (Brouhard et al., 2008). The variable C-terminal region in ch-TOG likely contains a cryptic sixth TOG domain and a further small helical domain (Hood et al., 2013; Burgess et al., 2015; Rostkova et al., 2018). TACC3 is a member of the transforming acidic coiled coil (TACC) family of proteins that have a long coiled-coil region in their C-terminus, which is expected to govern their homodimerization (Peset and Vernos, 2008). TACC3 is a substrate of Aurora-A kinase with phosphorylation of a residue in the ACID region permitting the interaction with clathrin (Burgess et al., 2018). The TACC3–ch-TOG interaction is evolutionarily conserved. Examples include: Alp7–Alp14 (yeast), TAC-1–Zyg9 (nematode), D-TACC–Msps (fly) and maskin–XMAP215 (frog). Mutational analysis has mapped the interaction between TACC3 and ch-TOG to a stutter in the TACC3 coiled coil (residues 678-688) and a folded region (residues 1932-1957; C-terminal to TOG6) in ch-TOG (Hood et al., 2013; Burgess et al., 2015; Gutiérrez-Caballero et al., 2015; Rostkova et al., 2018). However, the structural details of this interaction are not yet determined.

The interplay between ch-TOG and TACC3 at spindle poles is particularly unclear. In *Drosophila*, D-TACC concentrates Msps at the centrosome (Lee et al., 2001). In vertebrates, the interaction between TACC3 and ch-TOG is thought to recruit γ-tubulin ring complex (γ-TuRC) to the centrosome (Singh et al., 2014; Rajeev et al., 2023). XMAP215 can nucleate MT growth from MT seeds and γ-TuRCs, with the C-terminal region binding to γ-tubulin and N-terminal TOG domains stimulating nucleation (Thawani et al., 2018). Related to this, ch-TOG was shown to recruit γ-TuRC and activate it in interphase (Ali et al., 2023). Whether or not TACC3 is involved in this activity is unclear. In human cells, the localization of ch-TOG at mitotic centrosomes is independent of TACC3, which seems to be located distal to the centrosome (Gutiérrez-Caballero et al., 2015). In agreement with this, evidence from mammalian oocytes – where the spindle is formed without centrosomes – is that TACC3 forms a liquid-like domain at spindle poles (Fu et al., 2013; So et al., 2019). Finally, back to *Drosophila*, recent work indicates that D-TACC forms a liquid-like scaffold in the pericentriolar material (PCM) and not at the centrosome itself (Wong et al., 2024).

In this paper, we describe the structural details of the TACC3–ch-TOG interaction and report the discovery of Affimers that target the ch-TOG binding site on TACC3. The binding of Affimers occludes this site such that ch-TOG cannot associate with TACC3 in live cells. We use these inhibitory tools to demonstrate that the TACC3– ch-TOG interaction is required for PCM integrity during mitosis in human cells.

## Results

To produce a detailed model for the TACC3–ch-TOG interaction, biophysical and structural analysis of both proteins, alone and in complex, were carried out.

### TACC3 is a parallel coiled-coil dimer

To determine the oligomeric state of the TACC domain of TACC3, analytical ultracentrifugation (AUC) experiments were performed on TACC3 629-838. Sedimentation velocity traces were fitted well for all cells with one predominant species observed at 1.4 S. The frictional ratio was *∼*2.2 but varied depending on the prevalence of an additional small species which appeared to reduce the frictional ratio in proportion to its abundance (Supplementary Figure S1). The high frictional ratio is consistent with the interpretation that TACC3 629-838 is an extended molecule: a rod shape with potentially unfolded parts at either end. The molecular weight in solution, calculated from sedimentation coefficient and frictional ratio, is *∼*48–50 kDa indicating that the protein is dimeric. A very small amount of a heavier species was present in all samples, which might represent a tetramer, but as it displayed no concentration dependence, may be disulphide-crosslinked protein (Supplementary Figure S1).

Next, to ascertain the orientation of the dimeric TACC3 TACC domain, electron paramagnetic resonance (EPR) spectroscopy was performed. MTSL-labelling was carried on native cysteine residues in TACC3 629-838 C749A, C828A and TACC3 629-838 C662A, C749A mutants, generating TACC3 MTSL-C662 and TACC3 MTSL-C828, respectively. Continuous-wave EPR spectra of TACC3 MTSL-C662 and MTSL-C828 (Supplementary Figure S2A) showed a very slight broadening in comparison to one another, in support of a short interspin label distance at the upper limit of applicability between 1.6 and 1.9 nm (Banham et al., 2008). The corresponding DEER traces were weakly resolved and it was not possible to extract a reliable distance measurement (Supplementary Figure S2B,C), consistent with an interspin distance at the lower range of borderline region of applicability. The presence of a dipolar interaction, although not well-defined in these experiments, supports TACC3 being a parallel dimer, where we would expect the labels to be in close proximity. These conclusions are consistent with existing X-ray crystal structures of TACC domain fragment (TACC3 758-838, PDB 5LXN and 5LXO) in which this shorter region of the TACC domain is a dimeric, parallel coiled-coil.

### Structural characterization of the ch-TOG C-terminal domain

We have previously shown that the C-terminal domain of ch-TOG (residues 1517-1957) is sufficient for robust binding to the TACC3 TACC domain (Hood et al., 2013). Nuclear magnetic resonance (NMR) spectroscopy studies with the C-terminal domain (residues 1591-1941) from the *Drosophila* homolog of ch-TOG, minispindles (Msps), identified two independently mobile folded domains connected by a *∼*10 amino acid highly flexible linker. The N-terminal subdomain (residues 1591-1850) was characterized as an additional sixth TOG, and the C-terminal subdomain (residues 1860-1941) as an α-helix bundle (Hood et al., 2013; Burgess et al., 2015). The equivalent regions are retained in ch-TOG: the sixth TOG domain maps to residues 1517-1802 and the C-terminal α-helix bundle to residues 1817-1957.

In vitro co-precipitation studies with truncated ch-TOG constructs showed that the ch-TOG C-terminal α-helix bundle (1817-1957) is sufficient for binding to TACC3 (Figure 1A). We proceeded to determine the structure of ch-TOG 1817-1957 by NMR spectroscopy. Spectra collected from stable isotope-labeled ch-TOG protein allowed full backbone and sidechain assignment (BMRB access code: 27235) (Rostkova et al., 2018). Structure determination was carried out using a combination of experimental NMR data comprising dihedral angle constraints extracted from backbone (HN, HA, CA, CB, C’) chemical shifts using DANGLE (Cheung et al., 2010), distances from 15N and 13C resolved 3D NOESY-HSQC spectra and residual dipolar coupling constants (RDCs) (Table 1).

**Figure 1.**
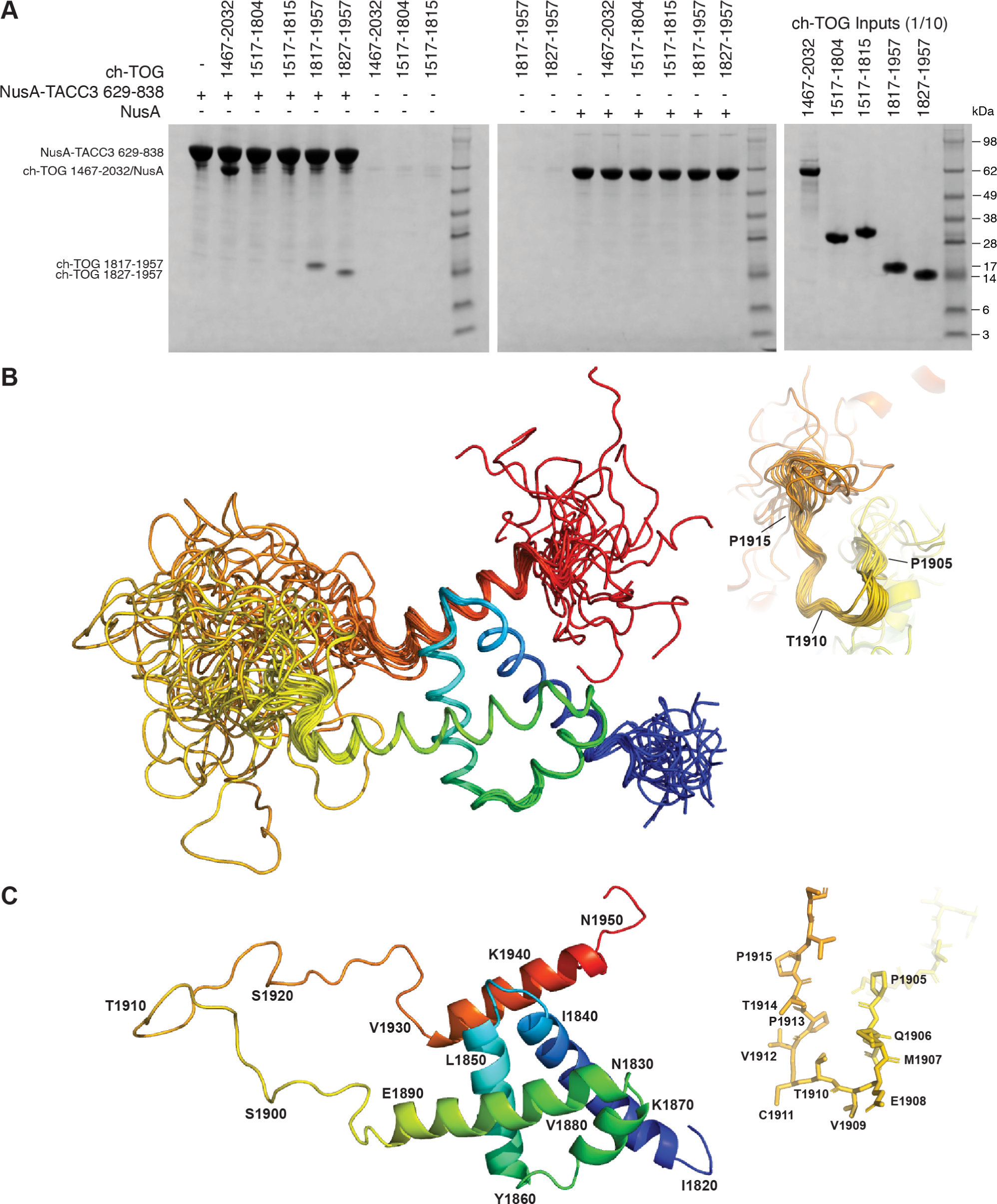
Characterization of ch-TOG 1817-1957. (**A**) In vitro co-precipitation assays between immobilized NusA-TACC3 TACC domain and ch-TOG truncates. (**B**) Superposition of the top 20 ch-TOG 1817-1957 NMR structures using the structured core of helices, H1-H4 (residues 1826-1894) for alignment. Inset, structures aligned on ch-TOG residues 1905-1915. (**C**) Cartoon representation of the best NMR structure for ch-TOG 1817-1957 shown in the same orientation as in (B). Inset, stick representation of ch-TOG residues 1905-1915. Structures in (B-C) are colored by spectrum mode where the N-terminus is blue and the C-terminus is red.

**Table 1.** Structure statistics and quality indices.

|  |  |
| --- | --- |
| Total NOEs | 3322 |
| H-bond constraints | 52 |
| Dihedral constraints | 174 |
| RDCs | 93 |

The best 20 NMR structures for ch-TOG 1817-1957 display good agreement within structurally ordered regions when superimposed (Figure 1B). The best individual structure for ch-TOG 1817-1957 is shown in Figure 1C. The domain consists of five helices, H1: 1825-1841; H2: 1846-1860; H3: 1865-1871; H4: 1874-1892 and H5: 1932-1945. Residues at the N- and C-terminus of the domain, 1817-1821 and 1946-1957, and the long linker region between H4 and H5, corresponding to residues 1898-1931, are disordered (Supplementary Figure S3).

The overall structure of the core domain (H1-H4) is composed of two pairs of V-shaped antiparallel α-helical hairpins connected by short loops. The V-shapes are created by the α-helices having residues with small side-chains proximal to the connecting loop, and larger side-chains nearer the other end. For the H1-H2 hairpin, these are G1849 at the narrow end, I1836 and I1853 in the middle and V1829, L1833 and Y1856 at the open end (Figure 2A). The H3-H4 hairpin has S1872 and S1873 near the loop and I1865, L1869 and L1884 near the open end (Figure 2B). These V-shaped hairpins are rotated by approximately 90° relative to each other and stacked flat on top of each other, exchanging hydrophobic contacts that hold the two hairpins together and form the core of the domain: H1 residues, V1829, L1833, I1836, F1837 and I1840; H2 residues, L1850, L1853, Y1854 and Y1856; H3 residues, I1865, F1868 and L1869 and H4 residues, F1876, V1880, L1884 and I1887. This hydrophobic core is devoid of polar amino acids and thus expected to be very stable (Figure 2C). By contrast, the interface with H5, which packs across the narrow end of the H1-H2 hairpin, is of more mixed character, consistent with unstable association; contacts are made between Y1935, L1939 and L1942 from H5 and K1839, E1835, K1838, S1842, N1845 and E1848 of the core domain (Figure 2D). The interaction may be further stabilized by the potential of salt bridges between residues R1938 & E1848, R1945 & E1844, R1943 & E1835, K1839 & E1835, E1855 & R1891, K1857 & E1881 and K1858 & E1888 (Figure 2). An intermediate structural state is observed for the short segment of residues P1905-P1915. This portion of the domain has very high RMSD values (Supplementary Figure S3). However, the distribution of ϕ/ψ values shows a more defined conformation (Supplementary Figure S3) with only modest variation; and superposing the structures using this region shows a good agreement of the structures (Figure 1B inset). This is supported by higher heteronuclear NOE values compared to the rest of the H4-H5 linker (Supplementary Figure S4). This portion of the domain also responds strongly to H5 binding to TACC3 (Figure 3A, see below).

**Figure 2.**
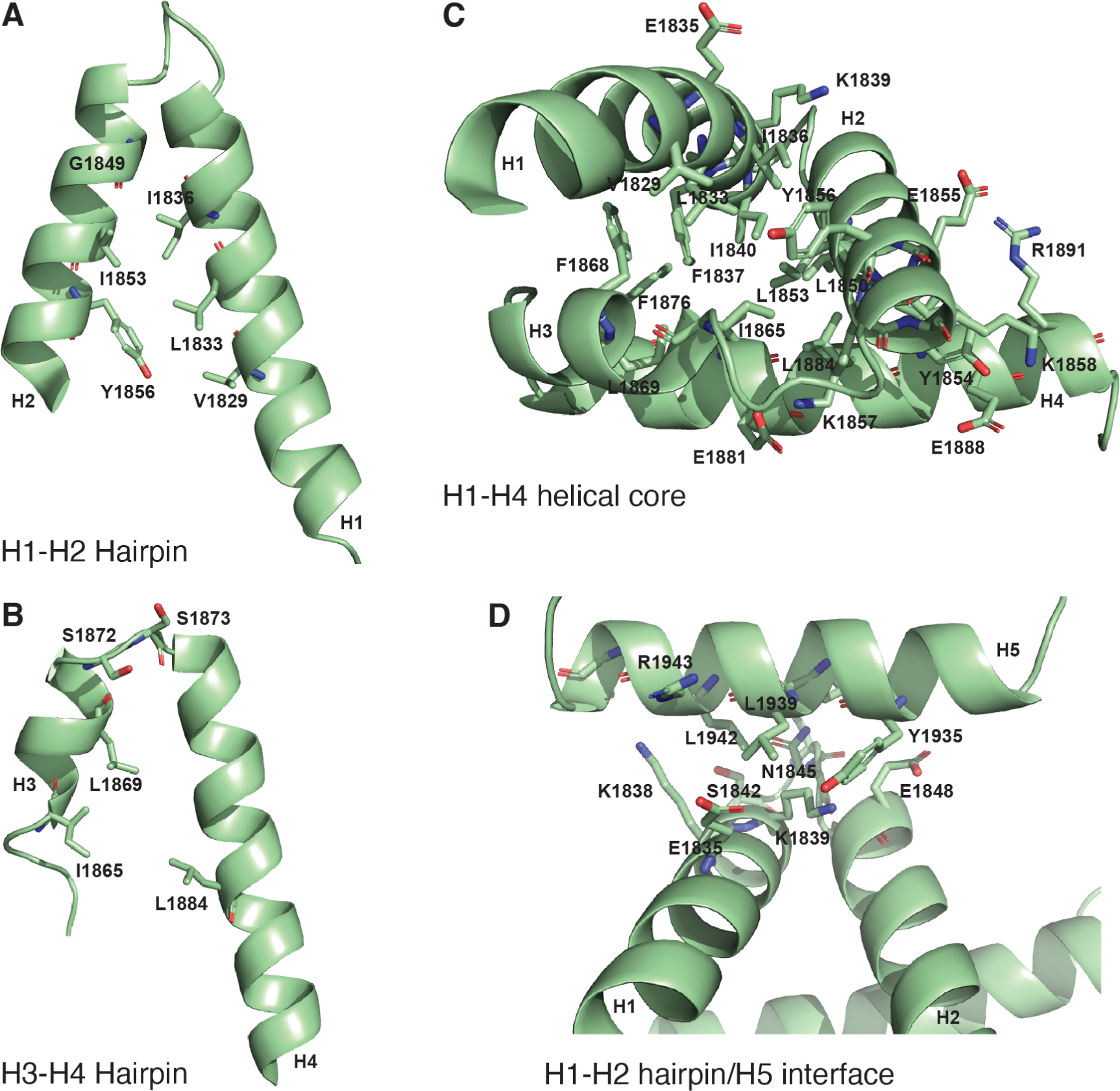
Structural interactions within ch-TOG 1817-1957. (**A**) Cartoon representation of the helical hairpin formed between H1 and H2. (**B**) Cartoon representation of the helical hairpin formed between H3 and H4. (**C**) Cartoon representation of the H1-H4 helical core. (**D**) Cartoon representation of H5 contacts with the H1-H2 hairpin.

**Figure 3.**
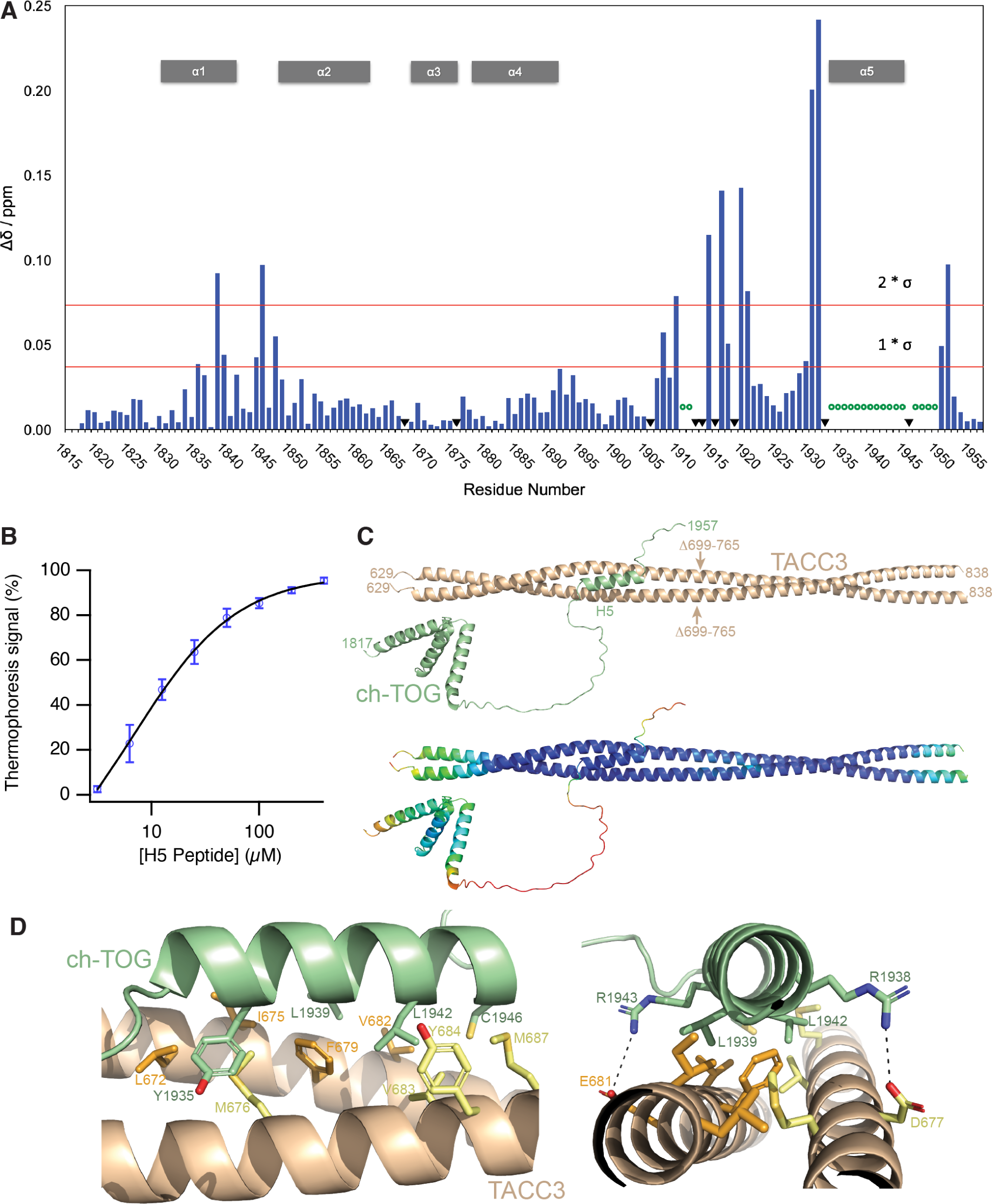
Interactions of ch-TOG with TACC3. (**A**) Chemical shift perturbations observed on interaction of ^2^H/^15^N labeled ch-TOG 1817-1957 with a three-fold excess of TACC3 629-838 Δ699-765. Data extracted from the spectra shown in Figure S5. (**B**) MST binding experiment of a synthetic ch-TOG H5 peptide with TACC3 629-838 Δ699-765. (**C**) Cartoon representation of an AlphaFold2 Multimer model of the complex between the TACC3 629-838 Δ699-765 dimer (wheat) and ch-TOG 1817-1957 (green). Arrows indicate the position of the deletion in TACC3. The model is shown below colored according to per residue confidence score (pLDDT) in rainbow colors from high (blue) to low (red) confidence. (**D**) Cartoon representations of the AlphaFold2 model showing the interface between the H5 region of ch-TOG (green) and TACC3 (wheat). Sidechains contributing to the interface from TACC3 protomer A (orange), TACC3 protomer B (yellow) and ch-TOG H5 (green), are shown in stick representation.

### Conformation and stability of ch-TOG 1817-1957

Residues in the core domain of the helix bundle (ch-TOG 1817-1957 H1-H4) have very low backbone RMSD values *∼*0.2 Å, large numbers of medium- and long-range NOEs, large positive heteronuclear NOEs, large ΔCA secondary chemical shifts and sizeable RDCs, indicating a tightly packed, well-folded and highly rigid region of the protein (Supplementary Figure S3 and S4). Within this region, only the loop region between H2 and H3 displays higher RMSD values (*∼*0.5 Å). Data corresponding to the N- and C-termini and the long H4-H5 linker (Figure 1B) are consistently in agreement with unstructured characteristics: large RMSDs, very few medium and long-range NOEs, negative or small heteronuclear NOEs, small CA secondary chemical shifts and very small RDCs (Supplementary Figure S3 and S4). H5 exhibits intermediate RMSD values of 0.3–0.4 Å (Supplementary Figure S3 and S4) and appears to be in an equilibrium between a stable α-helical state, in which it is associated with the rest of the domain, and a dissociated state in which it is partially unfolded.

### Interaction of ch-TOG 1817-1957 with TACC3

NMR titrations were performed with ^2^H/^15^N labeled ch-TOG 1817-1957 to which TACC3 629-838 Δ699-765 was added at ratio of 1:3, respectively (Supplementary Figure S5). All peaks corresponding to H5 disappeared upon binding to TACC3, with substantial chemical shift perturbations at positions neighboring H5 (Figure 3A). Modest chemical shift perturbations were also seen in the region around I1840, which is involved in the association of H5 with the H1-H4 core, and the short proline-rich segment in the middle of the linker that connects H4 to H5. These results indicate that only H5 (residues 1932-1945) of ch-TOG is involved in binding to TACC3. This is in agreement with previous work where deletion of ch-TOG 1932-1957 abolished binding to TACC3 (Gutiérrez-Caballero et al., 2015). As evidenced by the disappearance of H5 signals, while all other resonances remain visible, H5 is dislodged from the H1-H4 core concomitant with its binding to TACC3 (Figure 3A). Given the key role of ch-TOG H5 for complexation with TACC3, we tested whether a synthetic peptide corresponding to the H5 sequence retained the features required for interaction. Binding of the H5 peptide to TACC3 was measured with a *K*_D_ of 6.7 ± 1.5 µM (Figure 3B).

### Model of the TACC3–ch-TOG complex

A structural model of the interface between TACC3 and ch-TOG was generated by supplying AlphaFold2-Multimer (Evans et al., 2021; Jumper et al., 2021; Mirdita et al., 2022) with two copies of the sequence of TACC3 629-838 Δ699-765 and one copy of ch-TOG 1817-1957. Consistent with our AUC and EPR spectroscopy data, and with existing crystal structures, this region of TACC3 is predicted with high confidence as a dimeric parallel coiled-coil. The structure predicted for ch-TOG 1817-1957 is consistent with our NMR data (Figure 1B): a core bundle of four α-helices is formed by residues 1817-1892, and a fifth α-helix formed by residues 1931-1946 (H5) is separated from the bundle by a long, disordered region. Rather than folding back onto the bundle, H5 is confidently predicted to be flipped out and interacting with the TACC3 dimer (Figure 3C and S6A). Consistent with the NMR data (Figure 3A), the interface with TACC3 is formed exclusively by H5, the core α-helical bundle having no consistently or confidently predicted involvement. H5 binds into the groove between the two protomers of the TACC3 dimer in the region formed by residues 672-687. This binding site is consistent with previous observations that deletions of residues 678-681 or 682-688 of TACC3 abolished binding of ch-TOG (Hood et al., 2013). The interface creates a short stretch of trimeric coiled-coil, with the three amphipathic α-helices packing hydrophobic sidechains together to form its interior (Figure 3D). The principal hydrophobic sidechains contributed by ch-TOG are those of Leu1939 and Leu1942, which were previously identified as critical residues for the interaction (Gutiérrez-Caballero et al., 2015) (Supplementary Figure S6B), while Leu672, Ile675, Phe679 and Val682 are contributed by TACC3 protomer A, Met676 and Val683 by protomer B. Outside of the coiled-coil, intermolecular salt-bridges appear to be formed between Arg1943 of ch-TOG and Glu681 of TACC3 protomer A, and between Arg1938 of ch-TOG and Asp677 of protomer B. Mutation of ch-TOG Arg1938 and Arg1943 results in a reduction in binding between ch-TOG and TACC3 supporting the role of these residues in complex formation (Supplementary Figure S6B).

### Identification of TACC3 Affimers that inhibit the TACC3–ch-TOG interaction

To generate an inhibitor of the TACC3–ch-TOG interaction, we turned to Affimer technology, previously known as Adhiron technology (Tiede et al., 2014, 2017). Biopanning of an Affimer library against the C-terminal TACC domain of TACC3 (residues 629-838 Δ699-765) was carried out to identify antigen-specific clones. The subsequent clones were confirmed by phage ELISA and DNA sequencing, where they were grouped into six unique sequence families. A representative of each family was recombinantly expressed and purified for in vitro screening. Of the six Affimers, two precipitated heavily during purification and were not taken forward, while Affimers E4, E5, E7, and E8 were further characterized. *In vitro* co-precipitation assays using purified recombinant proteins, confirmed binding of Affimers E4, E7, E8 and E5 to the TACC3 TACC domain (Figure 4A). Upon addition of ch-TOG, formation of a ternary complex was observed in the case of Affimer E5, but not Affimers E4, E7 and E8, suggesting these bind at a site on the TACC domain that is required for ch-TOG association (Figure 4A). In ELISA experiments using purified proteins, Affimers E4, E7 and E8 exhibited clear binding to immobilized TACC3 TACC domain (residues 629-838 Δ699-765) (Figure 4B). Affimer E5 displayed high back-ground, indicating that it is prone to non-specific binding, and was not carried forward.

**Figure 4.**
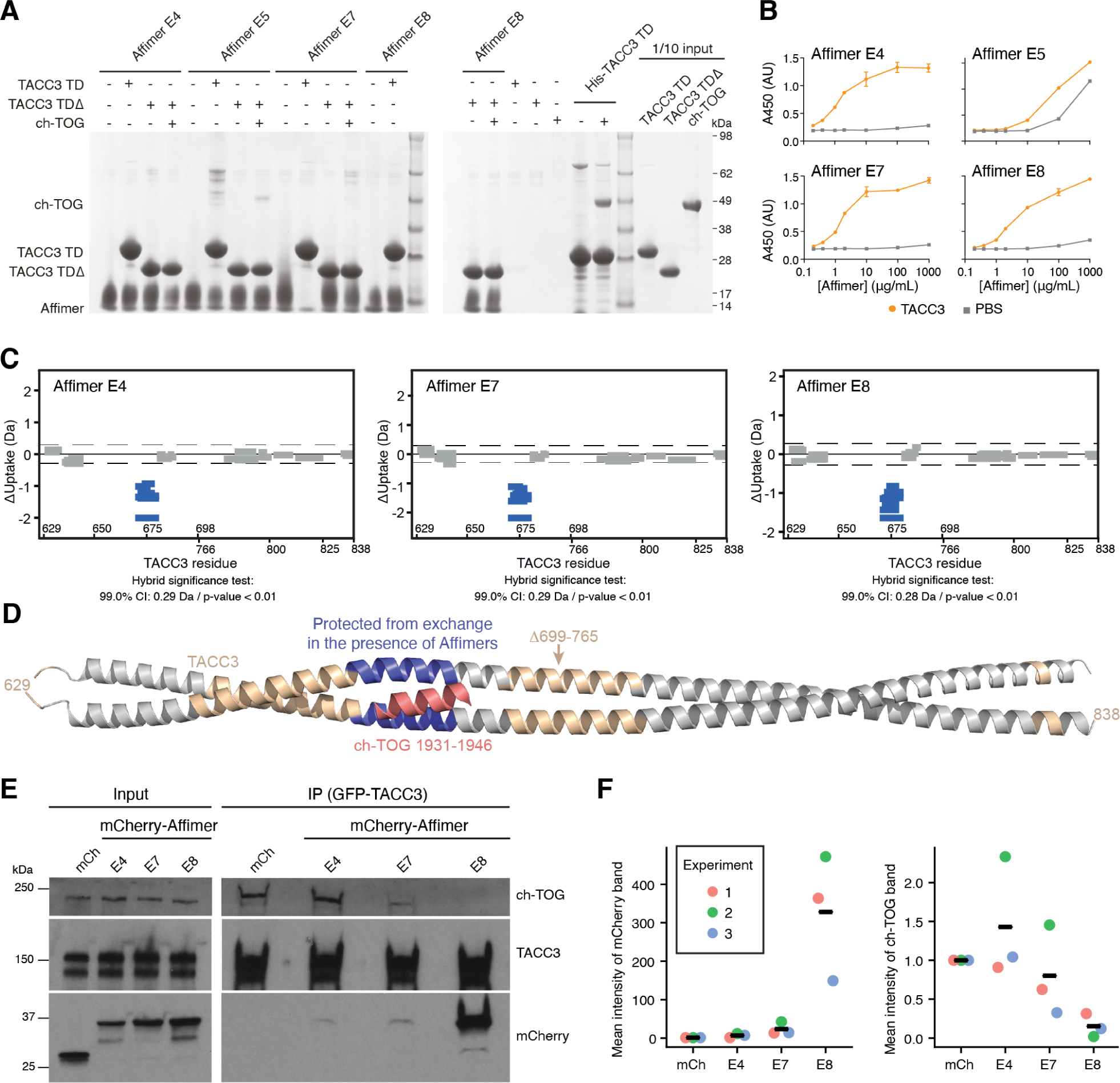
Isolation of Affimers that bind TACC3 and inhibit TACC3-ch-TOG interaction. (**A**) In vitro co-precipitation assay between Affimers, TACC3 and ch-TOG. C-terminal His-tagged Affimers were immobilized on nickel sepharose resin and incubated with TACC3 629-838 (TACC3 TD) or TACC3 629-838 Δ699-765 (TACC3 TDΔ). Binding of ch-TOG 1517-1957 in the presence of Affimer was assessed by the addition of ch-TOG to TACC3 TDΔ reactions. (**B**) ELISAs to assess binding between Affimers and TACC3 629-838 Δ699-765. Biotinylated TACC3 629-838 Δ699-765 was immobilized on Streptavidin coated plates and incubated with an Affimer dilution series (orange circles). Background binding of Affimers to the plate was measured by incubating the proteins in wells coated with PBS (gray squares). Data points are the mean ± standard error of the mean from two experiments. (**C**) Woods plots describing differences in deuterium uptake by residue, after 30 minutes of exchange, between TACC3 629-838 Δ699-765 in the absence of binding partner and in the presence of Affimers (as indicated). Woods plots were generated using Deuteros 2.0. Peptides colored in blue are protected from hydrogen/deuterium exchange in the presence of Affimers. Peptides exhibiting no significant difference in exchange between conditions, determined using a 99% confidence interval and a hybrid statistical test (dotted line), are shown in gray. (**D**) Cartoon representation the TACC3 629-838 Δ699-765–ch-TOG complex model with TACC3 colored according to HDX behavior as in (C). The region of TACC3 protected from hydrogen/deuterium exchange in the presence of Affimers E4, E7 and E8 is colored blue, and ch-TOG H5 is colored pink. (**E**) Pull-down assay using cell extracts from asynchronously growing HeLa cells expressing GFP-TACC3 and the indicated mCherry-Affimers, or mCherry as a control. GFP-TACC3 was immunoprecipitated using GFP-trap and a representative Western blot from three experiments is shown. Blots were probed for TACC3, mCherry and ch-TOG, as indicated. (**F**) Quantification of mCherry and ch-TOG bands from pull-down assays. Each dot represents the mean intensity of the indicated protein band normalized to the mCherry condition, colored by experiment. Crossbar indicates the mean from three experiments.

Next, we identified the binding sites of Affimers E4, E7 and E8 on TACC3, using hydrogen/deuterium exchange mass spectrometry (HDX-MS) (Figure 4C and Supplementary Figure S6C-F). The region of TACC3 protected from exchange by the binding of each of these Affimers was residues *∼*670 to 682, corresponding to the ch-TOG binding site (Figure 4D). This indicates that Affimers E4, E7 and E8 block the TACC3–ch-TOG interaction by direct occlusion of the binding site on TACC3. Having confirmed inhibition in vitro, we next wanted to test in human cells, the ability of the Affimers to bind TACC3 and to interfere with the TACC3–ch-TOG interaction. To do this, HeLa cells were transfected with plasmids expressing GFP-TACC3 and mCherry-tagged Affimers and GFP-trap was used to isolate GFP-TACC3 and any associated proteins from lysates. The immuno-precipitates were analyzed by Western blot, using antibodies to probe for TACC3, mCherry or ch-TOG (Figure 4E, F). Affimer E8 showed the strongest binding to GFP-TACC3 and an almost complete inhibition of the TACC3–ch-TOG interaction. Affimers E7 and E4 showed much weaker binding, with E7 but not E4 inhibiting the TACC3–ch-TOG interaction. None of the Affimers performed well as reagents for visualization or inducible relocalization of TACC3 (Supplementary Figure S7). However, we were able to see colocalization of each of the three Affimers with overexpressed GFP-TACC3 (Supplementary Figure S7). Together, these data suggest that in HeLa cells, Affimer E8 binds to TACC3 and inhibits an interaction with ch-TOG, whilst a smaller effect is observed with E7 and no effect is detected with E4, compared to the mCherry control.

### Disrupting the TACC3–ch-TOG interaction during mitosis

TACC3 and ch-TOG localize to the mitotic spindle in an Aurora-A dependent manner, where they are part of a multiprotein complex with clathrin and GTSE1 (TACC3–ch-TOG–clathrin–GTSE1) (Ryan et al., 2021). To test the effect of Affimers on subcellular localization of this complex, each Affimer was expressed in GFP-FKBP-TACC3 knock-in HeLa cells and the distribution of endogenous TACC3 and ch-TOG were compared at metaphase (Figure 5). As a positive control, cells not expressing Affimers were treated with the Aurora-A kinase inhibitor MLN8237 to abolish TACC3 and ch-TOG spindle localization, as described previously (Booth et al., 2011; Hood et al., 2013) (Figure 5A,C). A decrease in ch-TOG spindle localization comparable to that resulting from MLN8237-treatment was measured for cells expressing Affimers E7 and E8, with a more modest effect for E4 (Figure 5B,C). Surprisingly, a small decrease in spindle localization of TACC3 itself was also seen for all three Affimers with respect to untransfected cells (Figure 5B,C). However, the level of TACC3 at the spindle was three-fold higher than that in the cytoplasm and this was significantly higher than MLN8237-treated cells, suggesting that although there is a slight reduction, TACC3 remained enriched on the spindle. In an analogous set of experiments, we confirmed that the localization of endogenous clathrin at the mitotic spindle remained intact in CLTA-FKBP-GFP knock-in HeLa cells expressing TACC3 Affimers (Supplementary Figure S8). The complete disruption of ch-TOG localization in the presence of Affimers E7 and E8 is consistent with a model where ch-TOG requires an intact interaction with TACC3 for its localization to mitotic spindle microtubules (Hood et al., 2013; Ryan et al., 2021). Moreover, it suggests that the inhibition of the TACC3–ch-TOG interaction by the Affimers is specific and does not interfere with the other interactions within the TACC3–ch-TOG–clathrin–GTSE1 complex.

**Figure 5.**
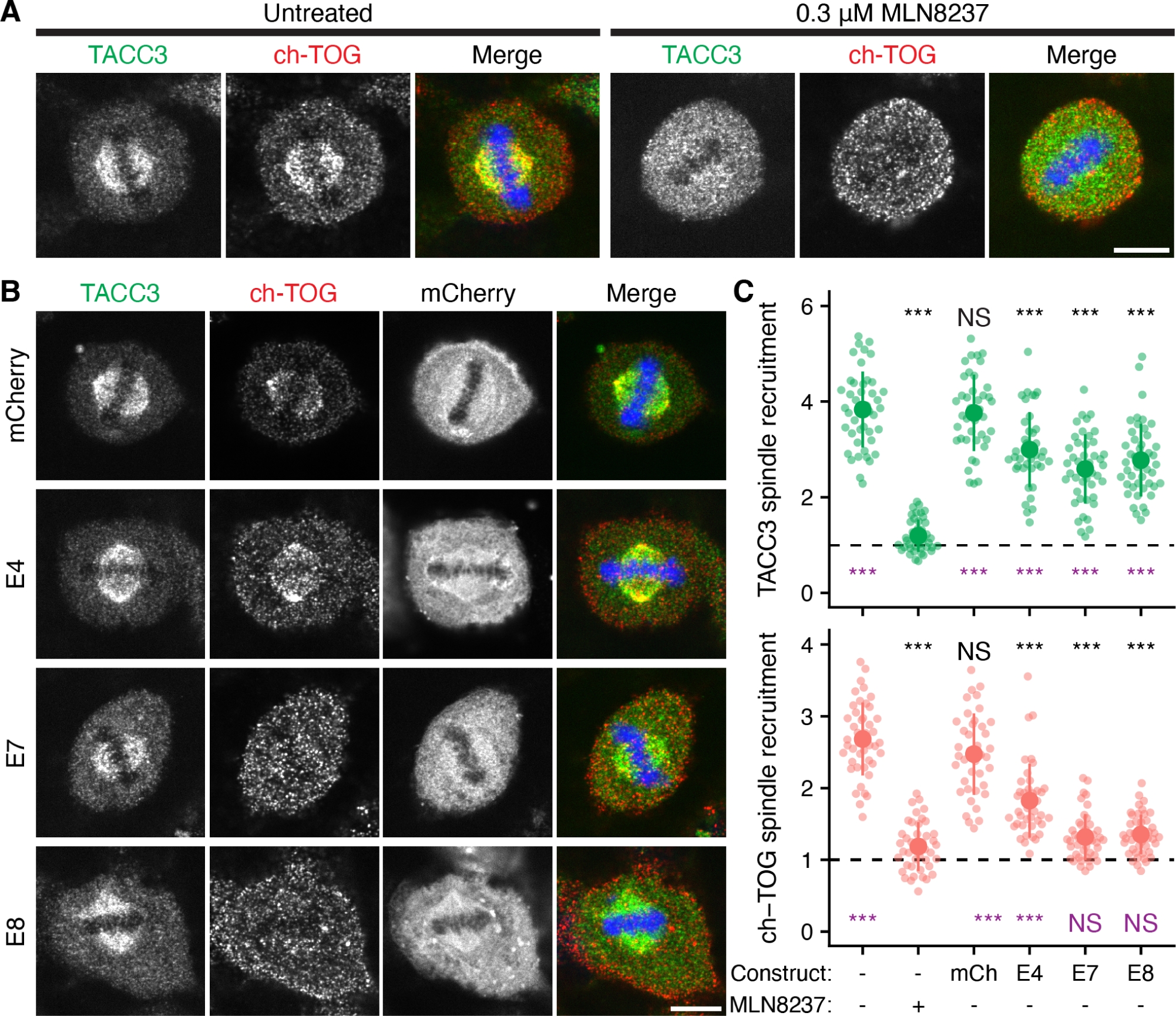
Affimer-mediated disruption of TACC3–ch-TOG interaction results in lower spindle localization of ch-TOG. (**A**) Representative confocal micrographs of untreated or MLN8237-treated (0.3 µM, 40 min) knock-in GFP-FKBP-TACC3 HeLa cells in metaphase. Cells were stained for ch-TOG (red), DNA (blue) and GFP-boost antibody was used to enhance the signal of GFP-FKBP-TACC3 (green). (**B**) Cells expressing mCherry or mCherry-Affimers, as labeled. Scale bars, 10 µm. (**C**) Quantification of spindle recruitment of TACC3 (green) and ch-TOG (red). Each dot represents a single cell, n = 40-45 cells per condition pooled from three independent experiments. Dashed line, no spindle enrichment. Large dot and bars, mean ± one standard deviation. Analysis of variance (ANOVA) followed by Tukey’s post hoc test is shown above each group, using the untransfected and untreated cells (black) and untransfected MLN8237-treated cells (purple) for comparison. ***, p < 0.001; NS, > 0.05.

### A role for the TACC3–ch-TOG interaction at the mitotic centrosome

Next we sought to ask whether targeted removal of ch-TOG from mitotic spindle microtubules would produce alterations in spindle morphology. HeLa cells expressing mCherry or mCherry-Affimers were fixed and stained for α-tubulin and pericentrin and imaged in 3D by confocal microscopy. Using a semiautomatic image processing pipeline, we measured several parameters including positioning, tilt, and scaling of the spindle and found no difference between cells expressing Affimers E7 or E8, compared with Affimer E4 or mCherry alone (Supplementary Figure S9). However, during this analysis, we noticed that many cells contained more than two distinct pericentrin foci, and that these additional foci were associated with small MT asters (Figure 6A). We therefore analyzed this using automated image analysis methods. A higher proportion of cells containing more than two pericentrin foci was observed in cells expressing the E7 (30.4 %) or E8 (33.3 %) Affimer, compared to E4 (8.2 %) or mCherry (2.2 %) (Figure 6B). Moreover, the additional pericentrin foci appeared smaller in size compared to the two foci that formed the bipolar spindle. Quantification of the total volume of pericentrin foci present in each cell revealed no significant difference in cells expressing Affimer E7 or E8 compared to the mCherry control (Figure 6C), suggesting that the additional foci are likely to represent fragments of the PCM, rather than amplified centrosomes. We concluded that, while the TACC3–ch-TOG interaction is not required for normal spindle morphology, it does appear to be required to maintain the structure of the pericentriolar material (PCM) during mitosis. To test this model, HeLa cells expressing mCherry or mCherry-Affimers were fixed and stained to visualize pericentrin and centrin1, as markers for the PCM and centrioles, respectively (Figure 6D). This analysis revealed that in the majority of cells containing >2 pericentrin foci, centrin-1 is absent from the additional sites (Figure 6E), confirming that they are detatched fragments of PCM.

**Figure 6.**
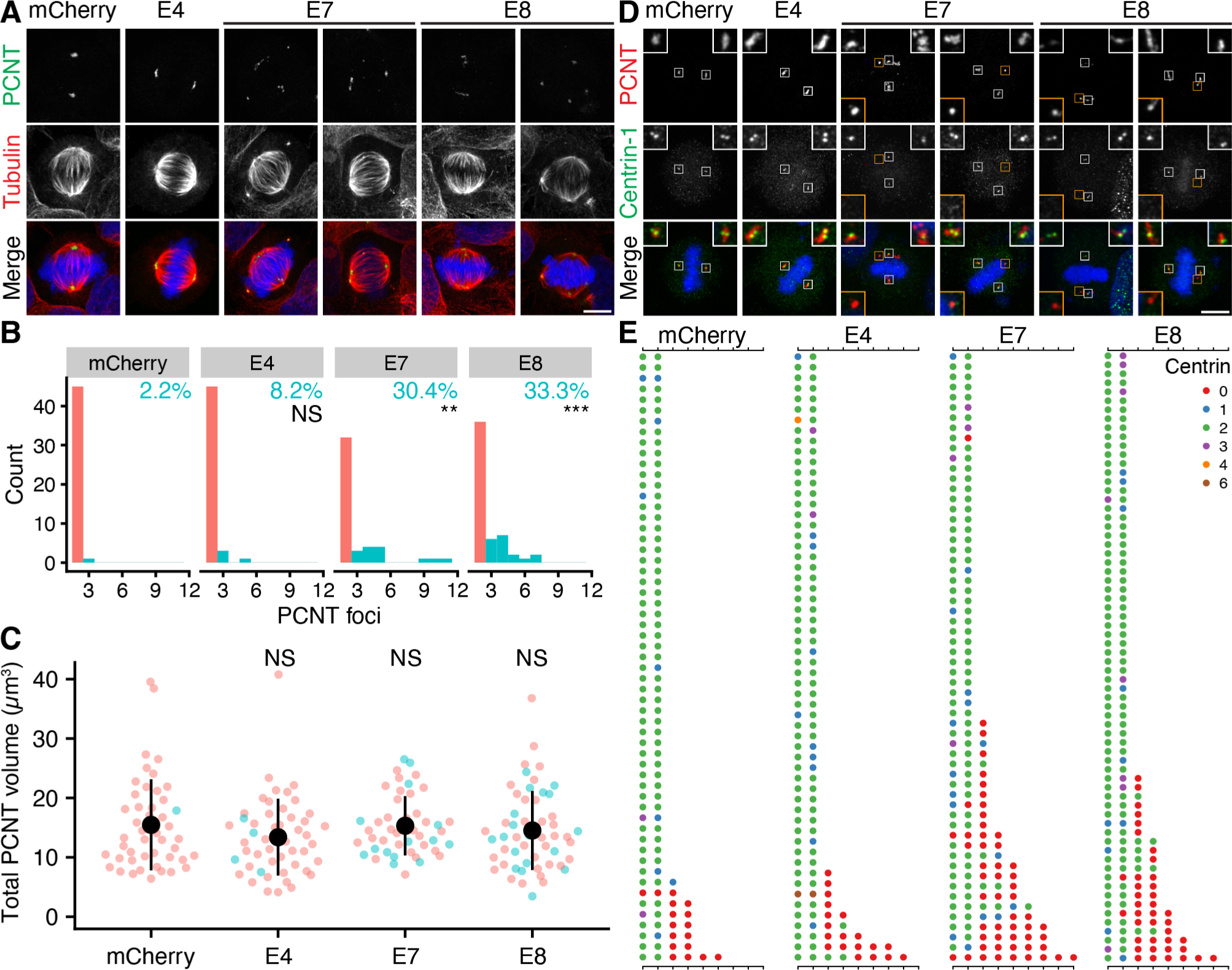
Expression of Affimers E7 and E8 leads to fragmentation of pericentrin. (**A**) Representative max intensity z projection images of cells expressing mCherry or mCherry-Affimers, as indicated; stained for pericentrin (PCNT, green), α-tubulin (red) and DNA (blue). (**B**) Histograms to show how many cells in each condition had 2 or more PCNT foci. The percentage of cells with >2 foci is indicated. Fisher’s exact test was used to test for association between the type of Affimer expressed and the PCM foci category. Bonferroni adjustment was used to calculate p values. (**C**) Dot plots show the total volume of pericentrin foci. Dots, single cells; n = 46-54 cells per condition over three independent experiments. Large dot and bars, mean ± one standard deviation. In B and C, cells with exactly 2 pericentrin foci are shown in salmon and those with >2, turquoise. Kruskal-Wallis test was used to compare the means between each group for the total pericentrin volume data. Dunn’s post hoc test with Bonferroni adjustment was used to calculate p values. The significance level is shown above each group, using the mCherry group for comparison. NS (not significant), p > 0.05; **, p < 0.01; ***, p < 0.001. (**D**) Representative max intensity z projection images of cells expressing mCherry or mCherry-Affimers, as indicated; stained for pericentrin (PCNT, red), centrin-1 (green) and DNA (blue). Insets show a 2.5 × zoom of the PCM, orange insets show sites of fragmentation. Scale bars, 10 µm. (**E**) Quantification of the number of pericentrin foci and the number of centrin-1 foci associated with each PCNT focus. Each dot represents a PCNT focus, with the color of the dot indicating the number of centrin-1 foci present, as described in the legend; each row is a different cell. Data is shown for n = 57-68 cells per condition over 3 independent experiments.

To address the possibility that Affimer E7- and E8-mediated PCM fragmentation might arise due to an off-target effect, we interfered with the TACC3–ch-TOG interaction by an alternative approach. In cells depleted of endogenous ch-TOG, we expressed RNAi resistant ch-TOG-GFP, a ch-TOG-GFP L1939A, L1942A mutant that cannot bind TACC3, or GFP as a control (Gutiérrez-Caballero et al., 2015). As expected, in ch-TOG-depleted cells expressing GFP, approximately 50 % of mitotic cells contained multipolar spindles (Figure 7A,B). This is consistent with previous reports of ch-TOG depletion, highlighting its role in spindle pole organization (Gergely et al., 2003; Cassimeris and Morabito, 2004). Expression of full-length ch-TOG rescued the multipolar phenotype, with only 14.2 % of cells containing >2 pericentrin foci (Figure 7A,B). In contrast, expression of the ch-TOG L1939A, L1942A mutant was associated with a pericentrin fragmentation phenotype in 36 % of cells (Figure 7A,B). These results were similar to those observed in the presence of Affimers E7 and E8 (Figure 6), and again the total volume of pericentrin in cells with fragmentation was similar to that in cells expressing full-length ch-TOG (Figure 7C). Taken together, these data suggest that the observed PCM fragmentation phenotype is a consequence of blocking the TACC3–ch-TOG interaction.

**Figure 7.**
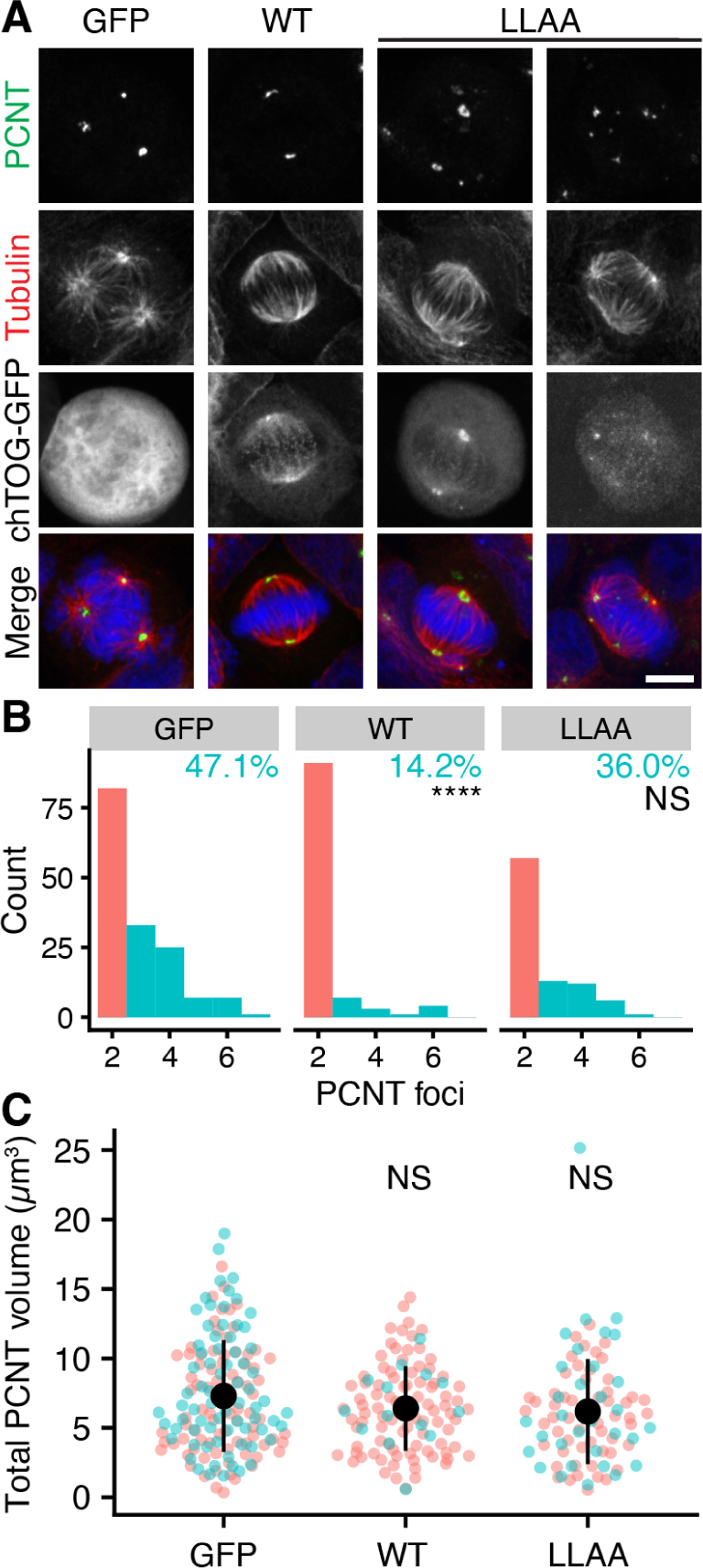
Expression of a ch-TOG mutant deficient in binding TACC3 results in fragmentation of pericentrin in mitotic HeLa cells. (**A**) Representative max intensity z projection images of HeLa cells coexpressing shRNA against ch-TOG, and either GFP, RNAi-resistant ch-TOG-GFP (WT) or ch-TOG(L1939,1942A)-GFP (LLAA). Cells were stained for pericentrin (green), α-tubulin (red) and DNA (blue). Scale bar, 10 µm. (**B**) Histograms to show how many cells in each condition had 2 or more PCNT foci. The percentage of cells with >2 foci is indicated. Fisher’s exact test was used to test for association between the protein expressed and the PCM foci category. Bonferroni adjustment was used to calculate p values. (**C**) Dot plots show the total volume of pericentrin foci. Dots, single cells; n = 89-155 cells per condition over three independent experiments. Large dot and bars, mean ± one standard deviation. In B and C, cells with exactly 2 pericentrin foci are shown in salmon and those with >2, turquoise. Kruskal-Wallis test was used to compare the means between each group for the total pericentrin volume data. Dunn’s post hoc test with Bonferroni adjustment was used to calculate p values. The significance level is shown above each group, using the GFP group for comparison. NS (not significant), p > 0.05; ****, p < 0.0001.

### TACC3–ch-TOG interaction is required for maintenance of centrosomal integrity during mitosis

From experiments in fixed cells, it was unclear whether fragmentation occurs prior to the cell entering mitosis, during spindle assembly, or once the spindle has formed. To clarify this point, we used live-cell imaging to visualize the PCM (mEmerald-γ-tubulin) and DNA (SiR-DNA) in cells expressing mCherry-Affimers, or mCherry as a control (Figure 8A). Consistent with our previous experiments, the proportion of cells with >2 γ-tubulin foci during metaphase, was higher in Affimer E7 (20.7 %) and Affimer E8 (19.4 %) expressing cells, compared to mCherry (6.7 %) and Affimer E4 (11.1 %). Interestingly, in cells expressing Affimers E7 and E8, the metaphase-anaphase transition was when the largest fraction of cells acquired supernumerary γ-tubulin foci, with 13.4 % and 15.3 % of cells displaying this phenotype, respectively, by this stage (Figure 8B). In these cells, the γ-tubulin foci underwent fragmentation to form smaller foci that remained close to the spindle before the cell divided (Figure 8A). Moreover, a prolonged metaphase-anaphase transition was observed in cells with this phenotype. Control cells maintained two distinct γ-tubulin foci throughout the observed stages and divided without a delay (Figure 8A). There was no significant change in overall mitotic progression in cells expressing Affimers E7 or E8 compared to Affimer E4 or mCherry alone (Figure 8C). However, when the mitotic timings of Affimer E7 and Affimer E8 expressing cells containing 2 or >2 γ-tubulin foci during metaphase were compared (Figure 8D), a delay in metaphase-anaphase progression was revealed: cells that exhibited PCM fragmentation during metaphase had a median metaphase-anaphase timing of 60 min (E7) and 102 min (E8), while those that cells that contained two foci throughout metaphase had a median metaphase-anaphase timing of 30 min. Taken together, these data show that inhibition of the TACC3– ch-TOG interaction causes fragmentation of the PCM during metaphase that is accompanied by a delay in the transition to anaphase.

**Figure 8.**
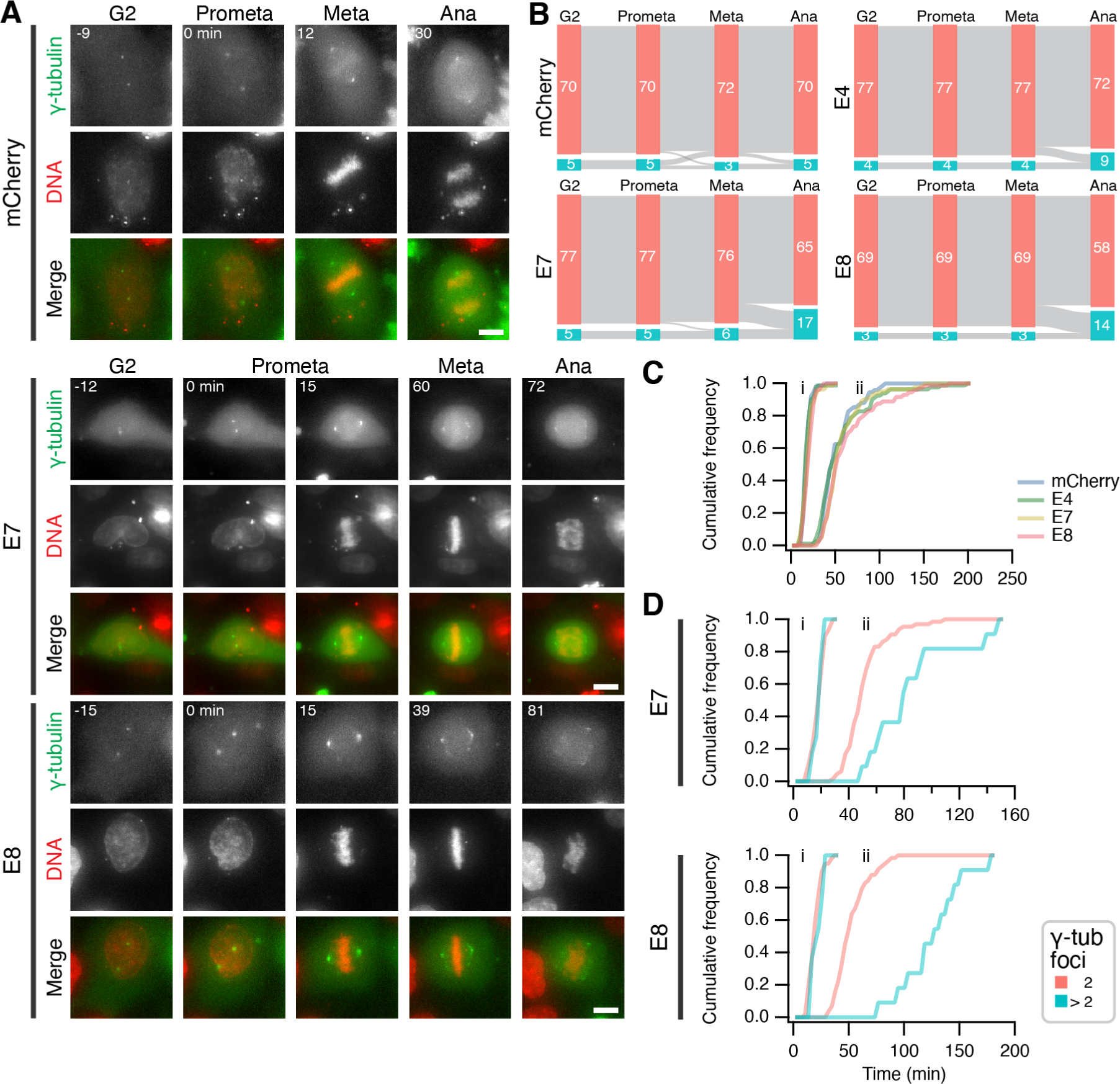
Blocking TACC3–ch-TOG interaction with Affimers results in fragmentation of PCM and mitotic delay. (**A**) Stills from live cell imaging experiments to track the number of γ-tubulin foci in cells expressing mEmerald-γ-tubulin (green) and the indicated mCherry or mCherry-Affimers constructs (not shown), SiR-DNA staining is shown (red). Scale bar, 10 µm. (**B**) Sankey diagrams to show the number cells containing supernumerary γ-tubulin foci at the indicated stages of cell division. The number of γ-tubulin foci was tracked from G2-prometaphase, prometaphase-metaphase and metaphase-anaphase. Numbers in each node represent the number of cells observed at each stage, as labeled. Node color represents the number of γ-tubulin foci in the cell, those with 2 are shown in salmon and those >2, turquoise. Data is pooled from four independent overnight experiments. (**C**) Mitotic progression of HeLa cells expressing mCherry or mCherry-Affimers. Cumulative histograms of prometaphase to metaphase (i) and prometaphase to anaphase (ii) timings. Number of cells analyzed: mCherry, 75; Affimer E4, 81; Affimer E7, 82; Affimer E8, 72. (**D**) Frequencies of Affimer E7- or Affimer E8-expressing cells shown in A, comparing timings of cells with two γ-tubulin foci (2; salmon) during metaphase with cells that undergo PCM fragmentation during metaphase (>2; turquoise). Number of cells: (2 and >2 foci, respectively): Affimer E7, 65 and 11; Affimer E8, 58 and 11.

## Discussion

In this paper, we described the structural details of the interaction between ch-TOG and TACC3. We isolated Affimers that bound TACC3 at the site of interaction with ch-TOG and displaced it. These new molecular tools could be expressed in cells and the function of the TACC3–ch-TOG interaction dissected. We uncovered a role for this interaction in stabilizing the pericentriolar matrix during mitosis.

The TACC3–ch-TOG interaction is through the H5 of ch-TOG binding to the parallel dimeric coiled-coil of TACC3 to form a short trimeric coiled-coil domain. In the absence of this interaction, H5 packs loosely against the four-helix core of this short domain and our NMR measurements showed the dynamicity of the interchange between these two states. The interior of the core region is entirely filled with hydrophobic amino acids and essentially all of them are highly conserved in the sequence of this domain from insects to mammals. This suggests that the core domain is a common feature of XMAP215/ch-TOG family proteins. The contacts made between H5 and the core domain are not strong, but they are sufficient to hold H5 in position and keep it in a helical conformation. Interestingly, in this position the two leucines (Leu1939 and Leu1942) that are important for binding to the TACC domain of TACC3 (Gutiérrez-Caballero et al., 2015) are pointing inwards and are not accessible from the outside. Therefore, in order to bind TACC3, H5 has to be dislodged from the core domain to make these leucines accessible. Such cryptic binding interfaces are seen in other proteins where a range of mechanisms are employed for their release (Gingras et al., 2006). How H5 becomes dislodged from the core in order to bind TACC3 and what triggers this event is an interesting question for future investigation.

The TACC3 Affimers developed in this study were sufficient to disrupt the TACC3–ch-TOG interaction in vitro and in cells. The specific targeting of a single protein-protein interaction is very useful, especially in the case of TACC3–ch-TOG where the proteins participate in other complexes, alone and in combination as well as likely functioning individually. Small molecule destabilization of TACC3, TACC3 depletion by RNAi, TACC3 knocksideways or Aurora-A inhibition all remove the entire TACC3–ch-TOG–clathrin–GTSE1 complex from the mitotic spindle as well as interfering with TACC3 functions at the spindle pole (LeRoy et al., 2007; Wurdak et al., 2010; Booth et al., 2011; Cheeseman et al., 2013; Hood et al., 2013; Akbulut et al., 2020). Similarly, depletion of ch-TOG has several other effects besides targeting the TACC3–ch-TOG interplay (Gergely et al., 2003; Herman et al., 2020). Targeting the interaction using Affimers means that TACC3 and clathrin stay in place on the mitotic spindle microtubules and the structure of the spindle is unaffected but that ch-TOG specifically is absent.

Interrupting the TACC3–ch-TOG interaction during mitosis in living cells resulted in a spindle pole defect: PCM fragmentation. TACC3 and ch-TOG have previously been shown to be involved in centrosome clustering in cancer cells; either via an interaction with integrin-linked kinase or with KIFC1 (Fielding et al., 2011; Saatci et al., 2023). However, the phenotype we uncovered is distinct since we saw multiple pericentrin foci in mitotic cells expressing the inhibitory Affimers, of which only two contained centrioles. Instead, the phenotype points to a recently described property of TACC3 at spindle poles. First in *Drosophila* spindles, the PCM is templated by a cnn-containing structure with a more liquid-like D-TACC region associated with it (Wong et al., 2024). Second in C. elegans oocyte meiotic spindles, XMAP215/ZYG-9 and TACC/TAC-1 act at multiple times during assembly to promote spindle pole integrity and stability (Harvey et al., 2023). Our data suggest that the TACC3–ch-TOG interaction is important for maintaining the PCM around the centrosomes and that fragmentation results in a mitotic delay. That the PCM fragmentation occurred midway through mitosis indicates that forces involved in chromosome congression likely contribute to the dispersal of PCM in the absence of the TACC3–ch-TOG interaction. The discovery of a novel mechanism involving TACC3–ch-TOG that maintains pericentriolar material integrity during mitosis underscores the importance of using precise tools such as Affimers to investigate protein-protein interactions.

## Methods

### Molecular biology and protein expression

The following plasmids were available from previous work: mNeonGreen-EB3, pMito-mCherry-FRBK70N, GFP-TACC3, pBrain-GFP-shch-TOG, pBrain-ch-TOGKDP-GFP-shch-TOG and pBrain-ch-TOGDPGFP(LL1939,1942A)-shch-TOG (Gutiérrez-Caballero et al., 2015); pETM6T1 TACC3 629-838, pETM6T1 TACC3 629-838 Δ699-765, pETM6T1 ch-TOG 1517-1957 and truncated ch-TOG constructs in pETM6T1 (Booth et al., 2011; Hood et al., 2013); mEmerald-γ-tubulin was from Addgene #54105. The expression construct for TACC3 629-838 Δ699-765-Avi was generated by adding a section encoding a non-cleavable C-terminal Avi-tag to the TACC3 629-838 Δ699-765 construct in pETM6T1 by primer extension PCR. TACC3 point mutations were introduced by QuikChange site-directed mutagenesis (Agilent). Selected Affimer cDNA sequences were subcloned from the phagemid display vector, pBSTG1 into pET11a (Tiede et al., 2014). All proteins were expressed and purified as described in earlier work (Hood et al., 2013). Co-precipitation assays performed with recombinant, purified proteins were also as described previously (Hood et al., 2013).

Sequences encoding TACC3-targeted Affimers E4, E7 and E8 were cloned into pmCherry-C1 and pmCherry-N1 vectors to give mCherry-Affimer or Affimer-mCherry constructs, respectively. The mCherry-Affimer orientation was found to be most effective.

### Analytical ultracentrifugation

Sedimentation velocity experiments were performed at 60k rpm in a Beckman XL-I analytical ultracentrifuge using the An-60 Ti rotor and Spin Analytical 2-sector cells. Samples of TACC3 629-838 were run at 2 mg mL^−1^, 0.75 mg mL^−1^ and 0.2 mg mL^−1^ in 50 mM Tris pH 7.5, 150 mM NaCl after extensive temperature equilibration at 4 °C and data collected in both absorbance and interferences modes. The vbar of the protein and the density and viscosity of the buffer at 4 °C were calculated using the program SEDNTERP (Philo, 2023). Sedimentation velocity traces were fitted with the c(s) with one discrete component model in SEDFIT. Parameters for the discrete component were fixed at their default values or left floating. The confidence level of maximum entropy regularization was varied between 0.68 and 0.95 to give essentially identical results.

### Electron Paramagnetic Resonance

MTSL labeled TACC3 proteins were produced as described in previous work (Concilio et al., 2016). Continuous-wave EPR (CW EPR) were measured to ascertain the possible presence of dipolar broadening. CW EPR spectra were recorded 120 K on a Bruker Micro EMX spectrometer using a super-high sensitivity probe head at 9.4 GHz using a microwave power of 20 mW and a modulation amplitude of 1 G. Pulsed electron-electron double resonance (PELDOR or DEER) spectroscopy measurements separates dipole-dipole coupling between spins, which is inversely proportional to the cube of their distance (Milov et al., 1981, 1984). It can measure distances between spin labels on the nanometer scale, typically between 1.5 and 6 nm (Jeschke, 2012). TACC3 629-838 C749A, C828A and TACC3 629-838 C662A, C749A (50 µM) containing 30 % glycerol were used for the studies. DEER measurements were performed at 50 K on a Bruker Elexsys 580 spectrometer. The four-pulse DEER sequence *π/*2*_νobs_ − τ*_1_ *− π_νobs_ − t − π_νpump_ −* (*τ*_1_ + *τ* 2 *− t*) *− π_νobs_ − τ*_2_ *− echo* was applied (Pannier et al., 2000), with *π/*2*_νobs_* pulse length of 16 ns, *π_νobs_* pulse length of 32 ns and *π_νpump_* pulse length of 32 ns. Pump pulses were applied at the maximum of the field sweep spectrum with the observe pulses 65 MHz lower. Phase-cycling was applied. The software, DEERAnalysis2022 (Jeschke et al., 2006), was used to subtract the exponential background decay due to intermolecular interactions and to calculate the interspin distance distribution by Tikhonov regularization.

### NMR spectroscopy

Protein expression, purification and NMR data collection was carried out as described previously (Rostkova et al., 2018). The assignment data yielded chemical shifts which were used by DANGLE (Cheung et al., 2010) to calculate backbone dihedral angle constraints. Distance constraints were extracted from 3D 15N and 13C resolved NOESY-HSQC spectra recorded at 800 MHz on a Bruker Neo spectrometer equipped with a TCI cryoprobe. Residual dipolar couplings were extracted from an In-Phase/Antiphase HSQC experiment (Ottiger et al., 1998) recorded in the presence of 5 mg/mL of Pf1 phage (ASLA biotech, Latvia). All NMR spectra processing was done in Topspin 3.4 (Bruker, Germany). All NMR data analysis and preparation for structure calculation was done using CCPNMR analysis 2.4 (Vranken et al., 2005; Skinner et al., 2015). Structure calculation was initiated with ARIA (Rieping et al., 2007) using the standard, default protocol. All NOESY distance constraints were initially unassigned allowing for the ARIA protocol to assign all distance constraints. Dihedral constraints were introduced in the 3rd round of structure calculation. Hydrogen bond constraints were introduced at the same time for Hydrogen bonds that were observed in at least 15 of the final 20 structures at the end of the previous calculation. Once the vast majority of distance constraints were assigned or an ambiguous assignment was confirmed after manual inspection, structure calculation was completed in XPLOR-NIH (Schwieters et al., 2003) at which point also the RDC constraints were introduced.

### Microscale Thermophoresis (MST)

The TACC domain (residues 629-838) and the synthetic H5 peptide (residues 1929-1947) were transferred into measurement buffer consisting of 20 mM sodium phosphate, 50 mM NaCl, 2 mM DTT, 0.05 % Tween and 0.02 % NaN_3_. The TACC3 domain was fluorescently labeled via NHS coupling using the standard NHS Red labeling kit following the manufacturers protocol (Nanotemper, Germany). MST experiments were recorded on a Nanotemper Monolith instrument using a concentration of the fluorescently labeled TACC domain of 600 nM while the concentration of the H5 peptide was varied in the range of 0–400 µM. Samples were measured using premium capillaries (Nanotemper, Germany). For the creation of the binding curve fluorescence intensity ratios were calculated from the reference reading prior to heating the sample to the end of the thermophoresis curve. Extracted fluorescence ratios were exported and analyzed in IgorPro 9 (WaveMetrics). The binding curve was fitted to a simple, two state equilibrium binding isotherm to obtain the affinity. The experiment was performed three times and the standard deviation of the three repeats was taken as the experimental error.

### Structural modeling

To generate models of TACC3 TACC domain in association with ch-TOG, the AlphaFold2 neural-network (Jumper et al., 2021) via the ColabFold pipeline (Mirdita et al., 2022) was used. The input proteins were based on fragments that interact biochemically: human TACC3 629-838 Δ699-765 (two copies) and human ch-TOG 1817-1957. ColabFold was executed using default settings: multiple sequence alignment with MMseqs2, no use of templates nor Amber, five models generated. Models were visualized using PyMol.

### Isolation of TACC3 Affimers

Screening was performed against biotinylated TACC3 629-838 Δ699-765-Avi as previously described (Tiede et al., 2014, 2017).

### ELISAs

All proteins were expressed and purified as previously described (Hood et al., 2013; Burgess et al., 2016). Biotinylated TACC3 Δ699-765 (50 µL of 10 µg mL^−1^, diluted in PBS) was immobilized on preblocked HBC Streptavidin plates (ThermoFisher) for 30 min at room temperature (RT). Negative control wells were generated by incubating wells with PBS instead of TACC3. Excess TACC3 was removed by washing wells 3 times with 300 µL PBS and 0.1 % TWEEN-20 (PBST). Affimers were diluted in PBS to generate a concentration series of 1 mg mL^−1^ to 0.2 µg mL^−1^ protein. 50 µL of each Affimer sample was applied to the wells and incubated for 1 h at RT. Plates were washed 3 times with PBST. 50 µL His-HRP antibody (Abcam ab1187, 1:5000) was diluted in PBS T20 Superblock (ThermoFisher), added to the wells and incubated for a further 30 min at RT. Plates were then washed 3 times with PBST. Binding was resolved by the addition of 50 µL TMB (ThermoFisher) and quenched by 50 µL 0.5 M sulphuric acid. Well absorbance was read at 450 nm.

### HDX-MS

HDX-MS experiments were conducted using an automated robot (LEAP Technologies) that was coupled to an Acquity M-Class LC with HDX manager (Waters). Samples contained 8.9 mM Na_2_HPO_4_, 1.5 mM KH_2_PO_4_, 137 mM NaCl, 2.7 mM KCl, pH 7.4. Experiments contained 8 µM TACC3 and 10 µM Affimer. To initiate the HDX reaction, 95 µL of deuterated buffer (8.9 mM Na_2_HPO_4_, 1.5 mM KH_2_PO_4_, 137 mM NaCl, 2.7 mM KCl, pD 7.4) was transferred to 5 µL of protein-containing solution, and the mixture was subsequently incubated at 4 °C for 0.5, 5 or 30 min. Three replicate measurements were performed for each time point and condition studied. 50 µL of quench buffer (8.9 mM Na_2_HPO_4_, 1.5 mM KH_2_PO_4_, 137 mM NaCl, 2.7 mM KCl, pH 1.8) was added to 50 µL of the labeling reaction to quench the reaction. Quenched sample (50 µL) was injected onto an immobilized pepsin column (Enzymate BEH, Waters) at 20 °C. A VanGuard Pre-column [Acquity UPLC BEH C18 (1.7 µm, 2.1 mm × 5 mm, Waters)] was used to trap the resultant peptides for 3 min. A C18 column (75 µm × 150 mm, Waters, UK) was used to separate the peptides, employing a gradient elution of 0–40 % (v/v) acetonitrile (0.1 % v/v formic acid) in H_2_O (0.3 % v/v formic acid) over 7 min at 40 µL min^−1^. The eluate from the column was infused into a Synapt G2Si mass spectrometer (Waters) that was operated in HDMSE mode. The peptides were separated by ion mobility prior to CID fragmentation in the transfer cell, to enable peptide identification. Deuterium uptake was quantified at the peptide level. Data analysis was performed using PLGS (v3.0.2) and DynamX (v3.0.0) (Waters). Search parameters in PLGS were: peptide and fragment tolerances = automatic, min fragment ion matches = 1, digest reagent = non-specific, false discovery rate = 4. Restrictions for peptides in DynamX were: minimum intensity = 1000, minimum products per amino acid = 0.3, max. sequence length = 25, max. ppm error = 5, file threshold = 3. Peptides with statistically significant changes in deuterium uptake were identified using the software Deuteros 2.0 (Lau et al., 2021). Deuteros was also used to prepare Woods plots. The raw HDX-MS data have been deposited to the ProteomeXchange Consortium via the PRIDE/partner repository with the dataset identifier, PXD052409 (Perez-Riverol et al., 2022). A summary of the HDX-MS data, as per recommended guidelines (Masson et al., 2019), is shown in Supplementary Table S1.

### Cell biology

HeLa cells (HPA/ECACC 93021013) or GFP-FKBPTACC3 knock-in HeLa cells (Ryan et al., 2021) were cultured in DMEM with GlutaMAX (Thermo Fisher) supplemented with 10 % FBS, and 100 U mL^−1^ penicillin/streptomycin. Cells were kept at 37 °C and 5 % CO_2_.

DNA transfection was by GeneJuice (Merck), according to the manufacturer’s instructions; using a 3:1 ratio of transfection reagent to DNA. Typically, cells were processed 48 h post-transfection. Aurora A inhibitor MLN8237 (Stratech Scientific) was used at 0.3 µM for 40 min. For induced relocation experiments, rapamycin (Alfa Aesar) was added to cells at a final concentration of 200 nM, 30 min prior to fixation.

### Immunoprecipitation

To isolate GFP-TACC3, HeLa cells expressing GFP-TACC3 were first lysed on ice for 30 min using lysis buffer (10 mM Tris/HCl pH 7.5, 150 mM NaCl, 0.5 mM EDTA, 0.5 % Nonidet™ P40 Substitute, 0.09 % sodium azide, protease and phosphatase inhibitor tablets). To pull down TACC3, 1.5 mg of lysate was incubated with GFP-trap (Chromotek) magnetic beads. To elute the target protein and any binding partners, beads were re-suspended in Laemmli buffer. Co-immunoprecipitation of the target protein(s) was assessed by SDS-PAGE followed by western blotting. The following primary antibodies were used: TACC3 (1:1000 Abcam ab134154); ch-TOG (1:1000 QED Bioscience 34032); mCherry (1:1000 Abcam ab167453). Followed by HRP-conjugated secondary antibodies (1:10,000) and detection by enhanced chemiluminescence.

To quantify the intensities of protein bands in Western blots, an ROI was manually placed on each band to be analyzed and the mean intensity was measured using Fiji. To correct for background noise, a ROI was placed above each band and the mean intensity was measured. The same size ROI was used for all measurements in a single experiment. Measurements were exported and read into R using a script where the mean pixel intensities were inverted and corrected for background noise. For each experiment, corrected values were normalized to the respective value from the mCherry control condition and plotted.

### Immunofluorescence

Cells were fixed with PTEMF (20 mM PIPES, pH 6.8, 10 mM EGTA, 1 mM MgCl_2_, 0.2 % Triton X100, and 4 % paraformaldehyde) for 10 min at room temperature, or with ice-cold methanol for 10 min. Following permeabilization in 0.5 % Triton X100 in PBS, cells were washed three times with PBS and then blocked at room temperature in 3 % BSA in PBS for 1 h, then incubated with primary antibodies diluted in blocking buffer for 1 h. Cover slips were washed three times with PBS three times, before incubation with AlexaFluor-conjugated secondary goat antibodies (Invitrogen) in blocking buffer for 1 h at room temperature. In experiments where CRISPR GFP-FKBP knock-in cell lines were used, anti-Rabbit GFP-boost (Invitrogen) or GFP-boost (Chromotek) antibodies were used to enhance the signal of GFP-tagged proteins. The following antibodies were used: Mouse anti-α-tubulin (1:1000 Sigma T6074); Rabbit anti-α-tubulin (1:2000 Invitrogen PA519489); Rabbit anti-ch-TOG (1:5000 QED Bioscience 34032); Rabbit anti-ch-TOG (1:800 Thermo PA5-59150); Rabbit anti-Pericentrin (1:5000 Abcam ab4448); Mouse anti-Centrin-1 (1:500 Sigma 04-1624); Mouse anti-TACC3 (1:1000 Abcam ab56595); anti-Rabbit-GFPboost (1:200 Invitrogen A-21311); GFP-boost (1:200 Chromotek gba488). Note that during this work, we evaluated the specificity of commercial ch-TOG antibodies for immunofluorescence (Shelford and Royle, 2020).

### Microscopy

Confocal imaging of fixed and live cells was performed using a Nikon CSU-W1 spinning disk inverted microscope equipped with a 2x Photometrics 95B Prime sCMOS camera using either a 100× oil (1.49 NA) or 60× oil (1.40 NA) objective, with respective pixel sizes of 0.110 µm) or 0.182 µm). Excitation was sequential, via 405 nm, 488 nm, 561 nm and 638 nm lasers, with 405/488/561/640 nm dichroic mirrors and Blue, 446/60; Green, 525/50; Red, 600/52; FRed, 708/75 emission filters. The system also contains an Okolab microscope incubator, Nikon motorized *xy* stage and a Nikon 200 µm *z*-piezo. Images were acquired with Nikon NIS-Elements software.

For mitotic progression experiments, the same microscope system was used, but in widefield mode. A 40× oil (1.30 NA) objective (pixel size, 0.28 µm) was used. Excitation was via a CoolLED (pE-300) light source, with Chroma ZET561/10× (mCherry), Chroma ZET488/10× (GFP) and Chroma ZT647rdc (FRed) excitation filters. Chroma ET575lp (mCherry), Chroma ET500lp (GFP) and Chroma ET665lp and Chroma ZT647rdc (FRed) dichroic mirrors were used with Chroma ET600/50m (mCherry), Chroma ET525/50m (GFP) and Chroma ET705/72m (FRed) emission filters.

For live-cell imaging experiments, cells were in 35 mm glass bottom fluorodishes with Leibovitz L15 CO_2_-independent medium supplemented with 10 % FBS at 37 °C.

### Image analysis

Analysis was done using Fiji, the data exported and read into R for further analysis and plotting. Spindle recruitment analysis was performed as described in (Ryan et al., 2021). Using a 1.4 µm^2^ ROI manually placed to measure the average fluorescence intensities of three regions of the spindle (away from the poles), the cytoplasm and one region outside of the cell as background. In R, following background subtraction, the average spindle value was divided by the average cytoplasm value to generate a spindle enrichment ratio and plotted.

To measure spindle morphology and positioning in 3D, a semi-automated procedure was used. First, image stacks were segmented (mCherry channel) using LabKit to obtain a segmented cell volume. The position of centrosomes, a line through the metaphase plate and markers to delineate the cell of interest were recorded and then, using LimeSeg, a skeleton of the cell was found. In IgorPro, all results were read in and spatial statistics were calculated. A sphere that best fit the surfels generated by LimeSeg was used to calculate the distances from each spindle pole to the cell boundary and to generate the spindle offset measurement. Note that, similar results were obtained in 2D using a routine written in R.

For the quantification of pericentrin and γ-tubulin foci, image stacks were analyzed using 3D Objects Counter in Fiji, and subsequent plotting R. Due to the small size of the centrin-1 foci and the tendency for overlap, 3D Objects Counter could not be used and so the counts were done manually by an experimenter blind to the conditions of the experiment. For mitotic progression experiments, the transition stages and number of γ-tubulin foci were recorded manually and analyzed in R (Sankey diagram) or IgorPro (Mitotic Timing).

### Statistical analysis

To compare among three or more groups, a one-way ANOVA was used with Tukey’s post hoc test. A Kruskal-Wallis test with Dunn’s post hoc test was used for data that did not follow a normal distribution. To assess normality, a Shapiro-Wilk test was used. Fisher’s exact test was used to test for an association between Affimer expression and PCM fragmentation. The Bonferroni correction method was used to adjust p-values to account for multiple comparisons.

### Data and software availability

The NMR co-ordinates of ch-TOG 1817-1957 have been deposited in the RCSB PDB and BMRB databases, with the identifiers PDB: 9F4C; BMRB: 34916. The mass spectrometry proteomics data have been deposited to the ProteomeXchange Consortium via the PRIDE partner repository with the dataset identifier PXD052409. All code used in the manuscript is available at https://github.com/quantixed/p062p035.

## ACKNOWLEDGEMENTS

EPR experiments were carried out at the EPSRC National Research Facility (NS/A000055/1). NMR experiments were recorded at the KCL Centre for Biomolecular Spectroscopy with the help of RA Atkinson. We thank Claire Mitchell and Laura Cooper from the Computing and Advanced Microscopy Unit (CAMDU) for their help and support. The work was supported by a Programme Award from Cancer Research UK (C25425/A27718) and a Project Award from BBSRC (BB/L023113/1). J.S. was supported by a studentship from the Biotechnology and Biological Sciences Research Council (BBSRC) Midlands Integrative Biosciences Training Partnership (MIBTP) (BB/M01116X/1). A.N.C. acknowledges support of a Sir Henry Dale Fellowship jointly funded by the Wellcome Trust and the Royal Society (grant number 220628/Z/20/Z). Funding from the BBSRC (BB/M012573/1) enabled the purchase of HDX-MS equipment.

## AUTHOR CONTRIBUTIONS

Author contributions are shown in Supplementary Figure S10.

## COMPETING FINANCIAL INTERESTS

Affimer reagents were invented by DCT and patented by the University of Leeds. The patent was licensed to Avacta Life Sciences, royalties are received as part of this agreement and it is managed by University of Leeds Commercialization Services.

## Supplementary Information

**Figure S1.**
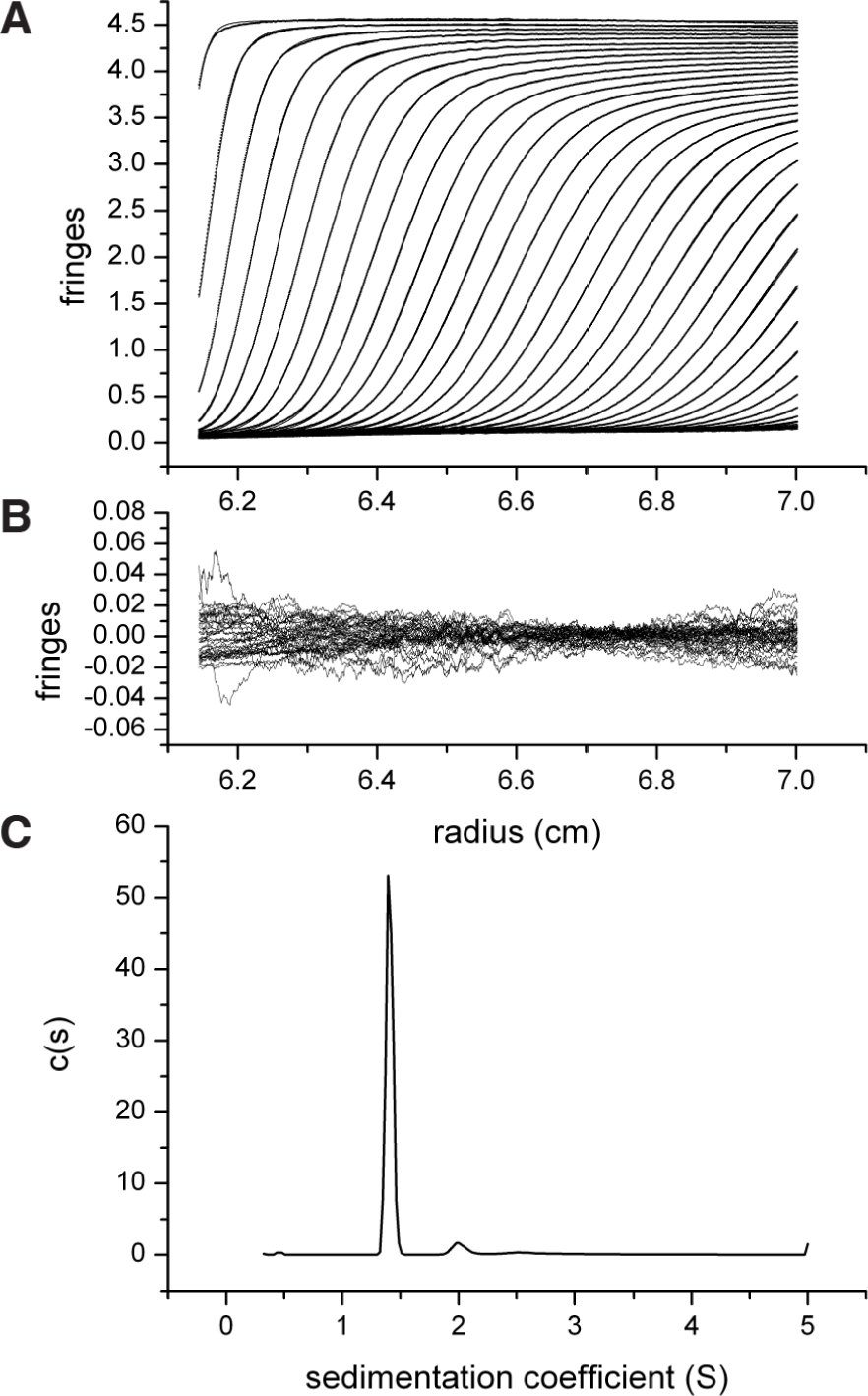
Sedimentation velocity traces and fits of TACC3 629-838 (2 mg/ml) (**A**) Fitting model: c(s)+1 discrete component, ME regularization at a confidence level of 0.68. (**B**) Residuals of fit shown in (A). (**C**) Resulting c(s) distribution.

**Figure S2.**
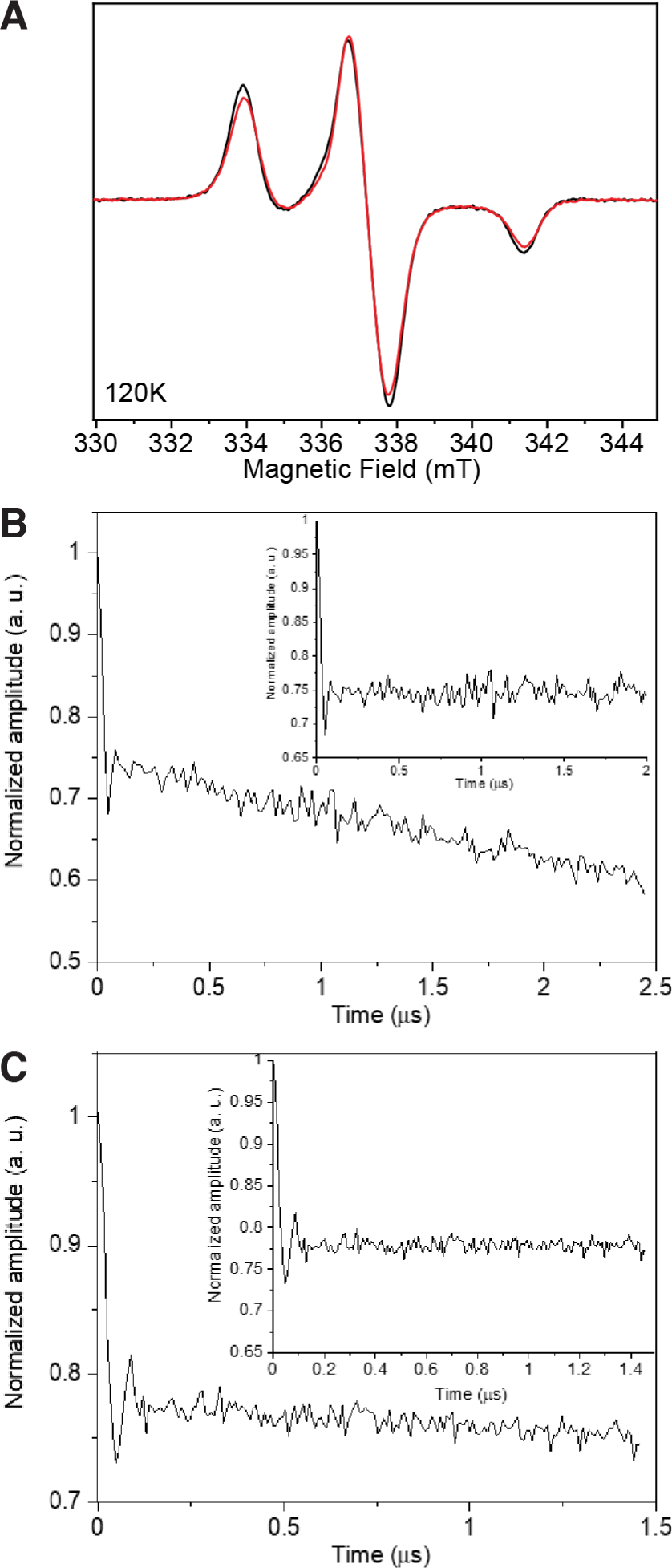
CW EPR and DEER of TACC3 TACC domain. (**A**) CW EPR spectra of TACC3 MTSL-C828 (black) and TACC3 MTSL-C662 (red) at 120 K. (**B-C**) Normalized four-pulse DEER trace at 50 K for (B) TACC3 MTSL-C828 and (C) TACC3 MTSL-C662. Inset, traces after subtraction of a mono-exponential decay. The weak oscillation is most likely from residual proton modulation.

**Figure S3.**
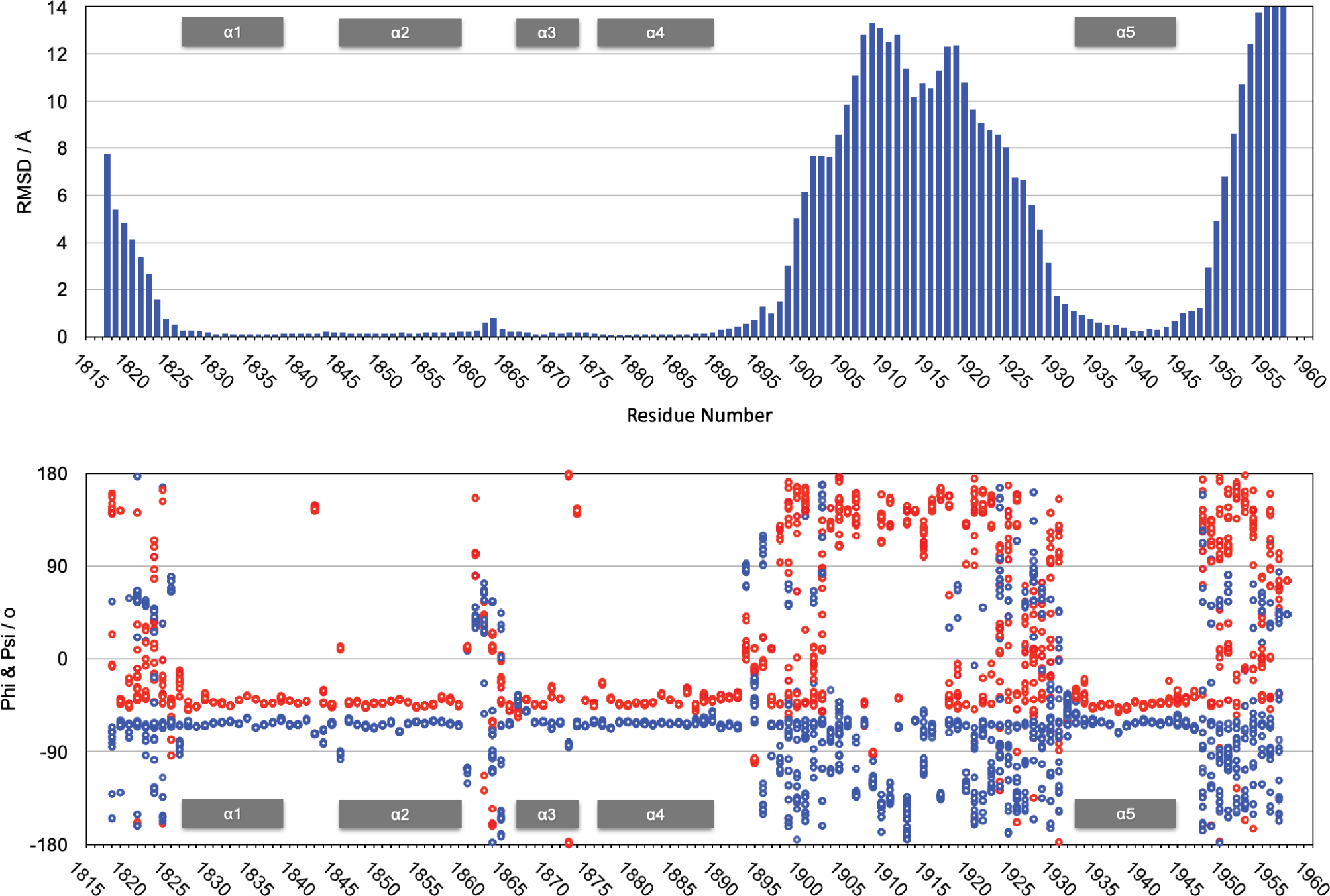
Variability plot of the top 20 ch-TOG 1817-1957 structures. Plots were based on backbone RMSD values (top) and a distribution of ϕ (blue) and ψ angles (red) (bottom) against amino acid sequence.

**Figure S4.**
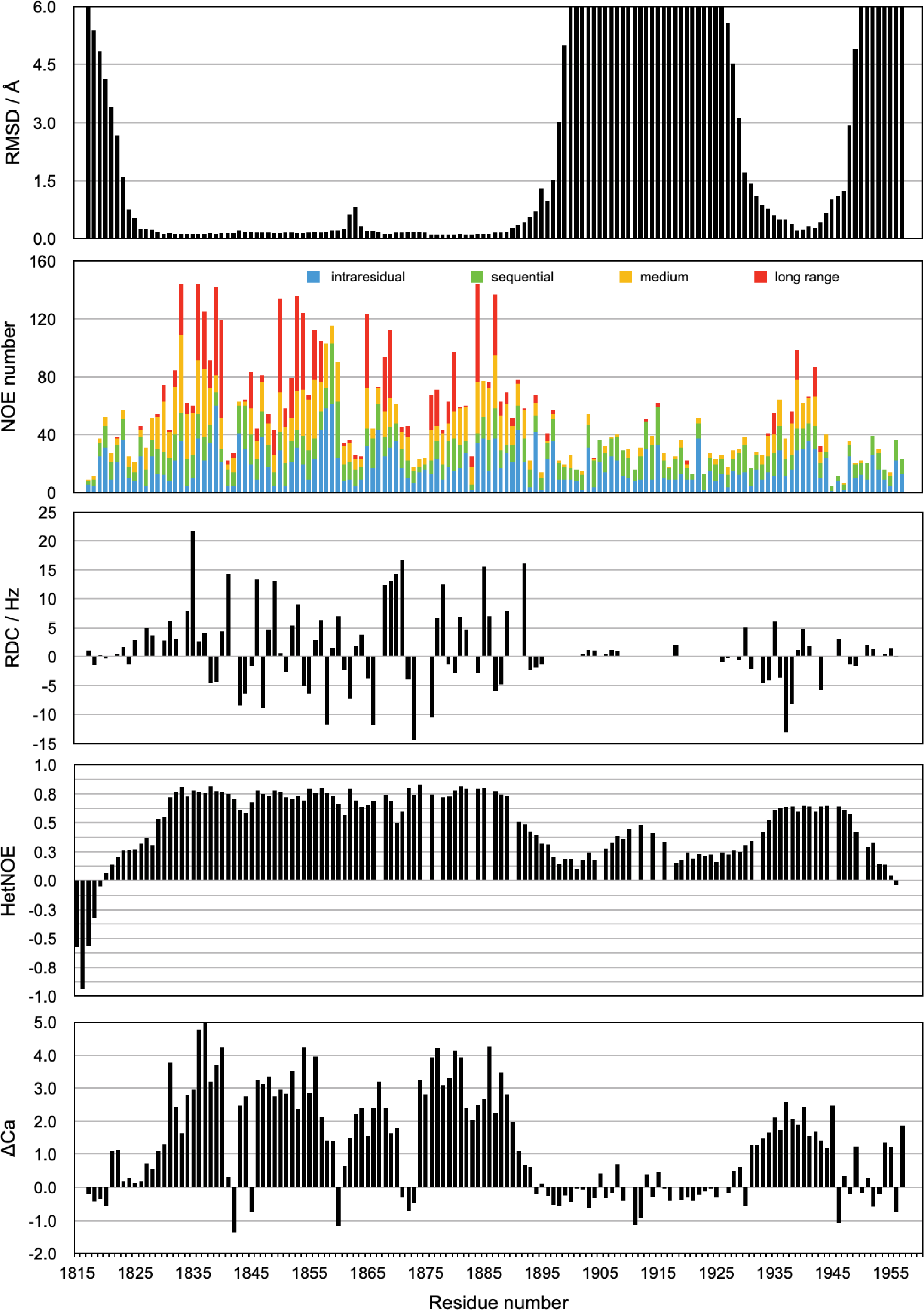
Conformation and stability of ch-TOG 1817-1957. The panels are from top to bottom: Backbone RMSD values (as shown in Figure S3), number of NOE derived distance constraints, backbone amide RDC values, heteronuclear NOE and Cα secondary chemical shifts, taken from Rostkova et al (2018).

**Figure S5.**
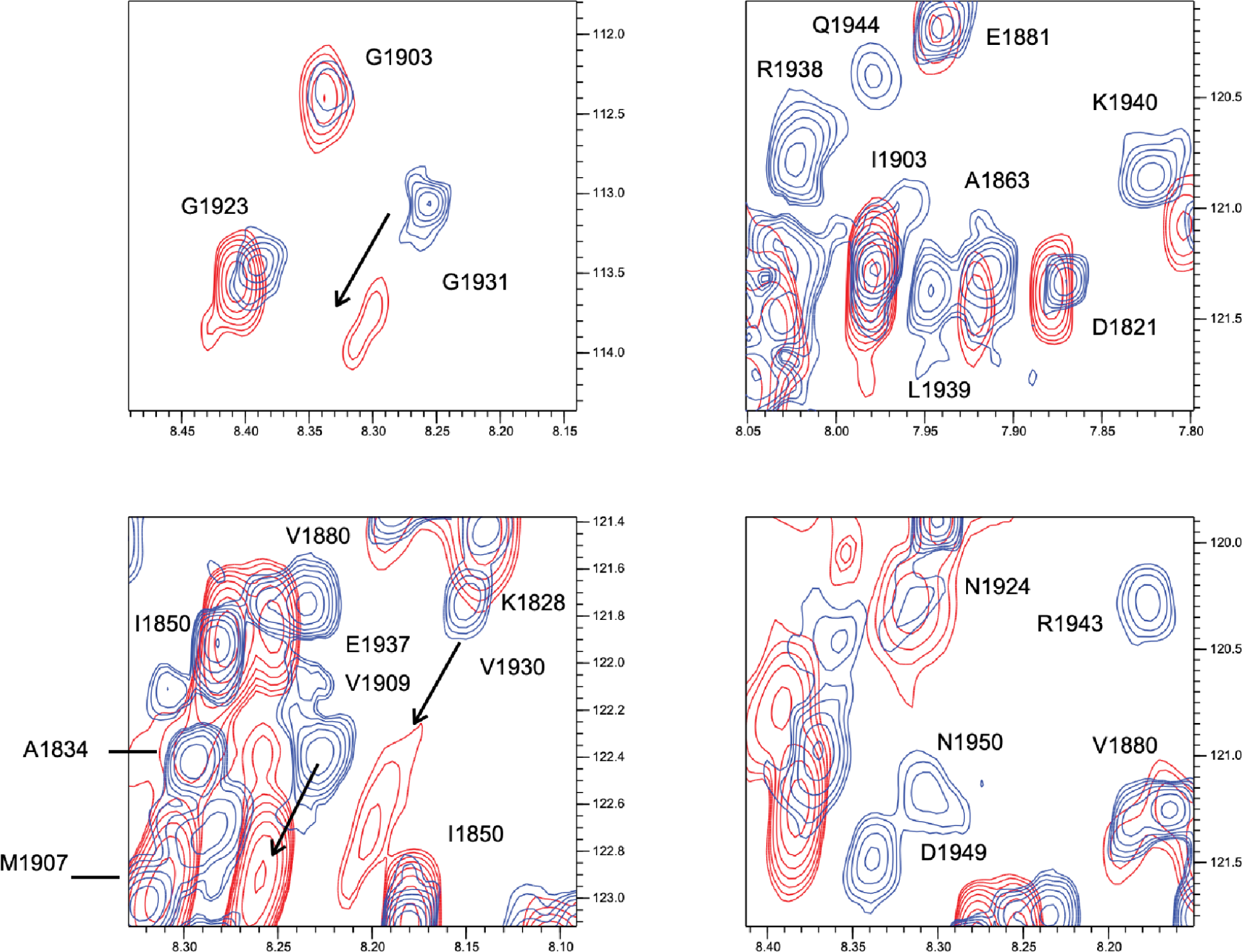
Portions of a TROSY experiment of 2H/15N labeled ch-TOG 1817-1957. Plots show the protein alone (blue) and in the presence of a three-fold excess of TACC3 629-838 Δ699-765 (red).

**Figure S6.**
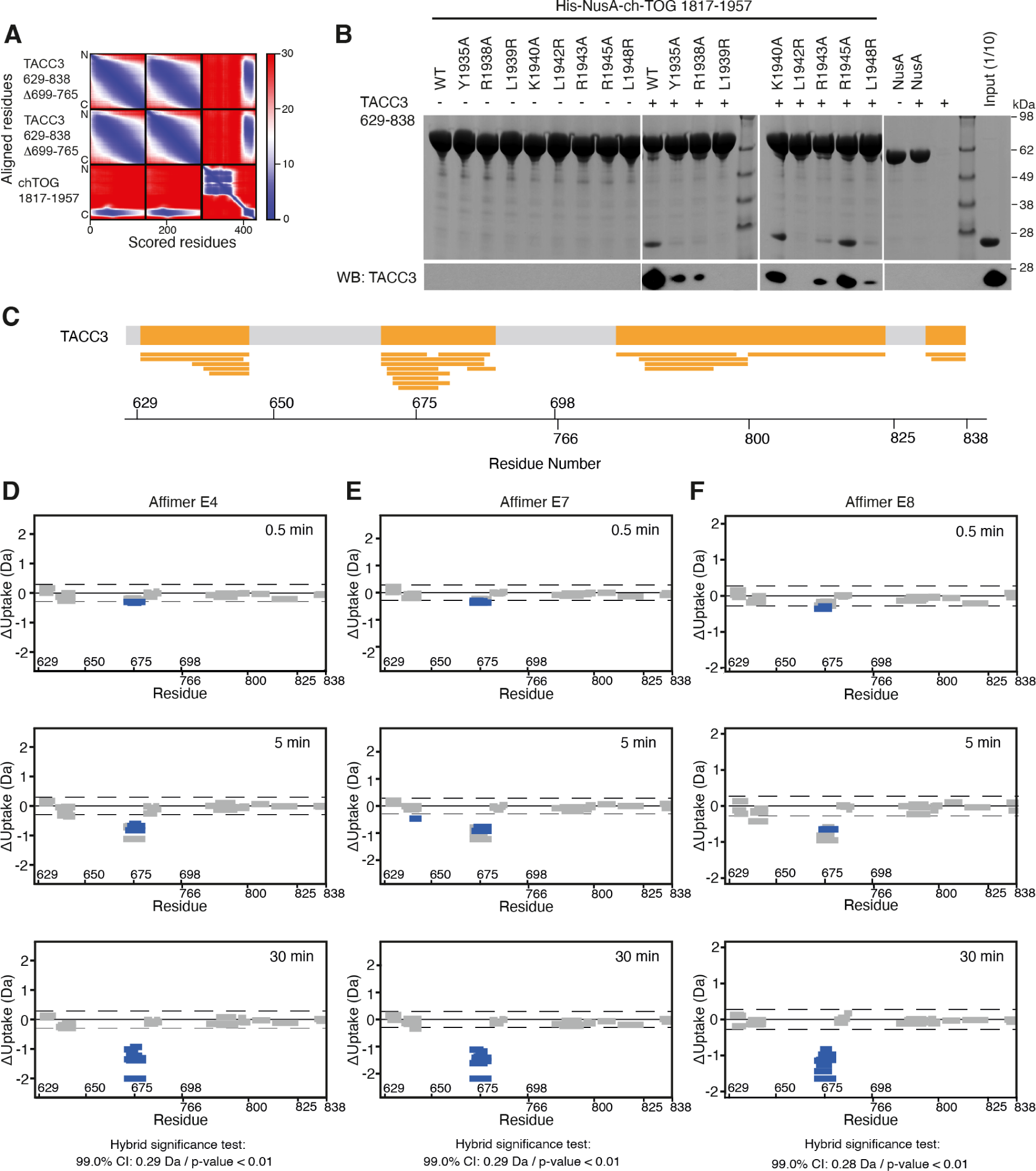
AlphaFold2 modeling of TACC3 and ch-TOG, the effect of ch-TOG H5 mutations on TACC3 binding and HDX-MS of TACC3 in the absence/presence of Affimers. (**A**) Predicted aligned error (PAE) plot for a model of the complex between TACC3 629-838 Δ699-765 and ch-TOG 1817-1957 generated by AlphaFold2 Multimer. (**B**) In vitro co-precipitation assays between immobilized His-NusA-ch-TOG 1817-1957 constructs and TACC3 629-838 (top). Binding of TACC3 was resolved by Western blot (bottom). (**C**) Sequence coverage map of TACC3 629-838 Δ699-765 in HDX-MS experiments. The yellow shaded regions in the thick bar at the top of the panel represent regions with sequence coverage while gray indicates regions that were not covered by detected peptides. Narrow yellow bars represent the individual peptides detected. (**D-F**) Woods plots showing the differences in deuterium uptake in TACC3 at three HDX timepoints (0.5, 5, 30 min of HDX), comparing TACC3 629-838 Δ699-765 alone with TACC3 629-838 Δ699-765 in the presence of Affimers E4 (D), E7 (E) and E8 (F). Woods plots were generated using Deuteros 2.0. Peptides colored in blue are protected from hydrogen/deuterium exchange in the presence of Affimers. Peptides with no significant difference in exchange between conditions, determined using a 99 % confidence interval and a hybrid statistical test (dotted line), are shown in gray. A summary of key details of the HDX-MS experiment is shown in Supplementary Table 1.

**Figure S7.**
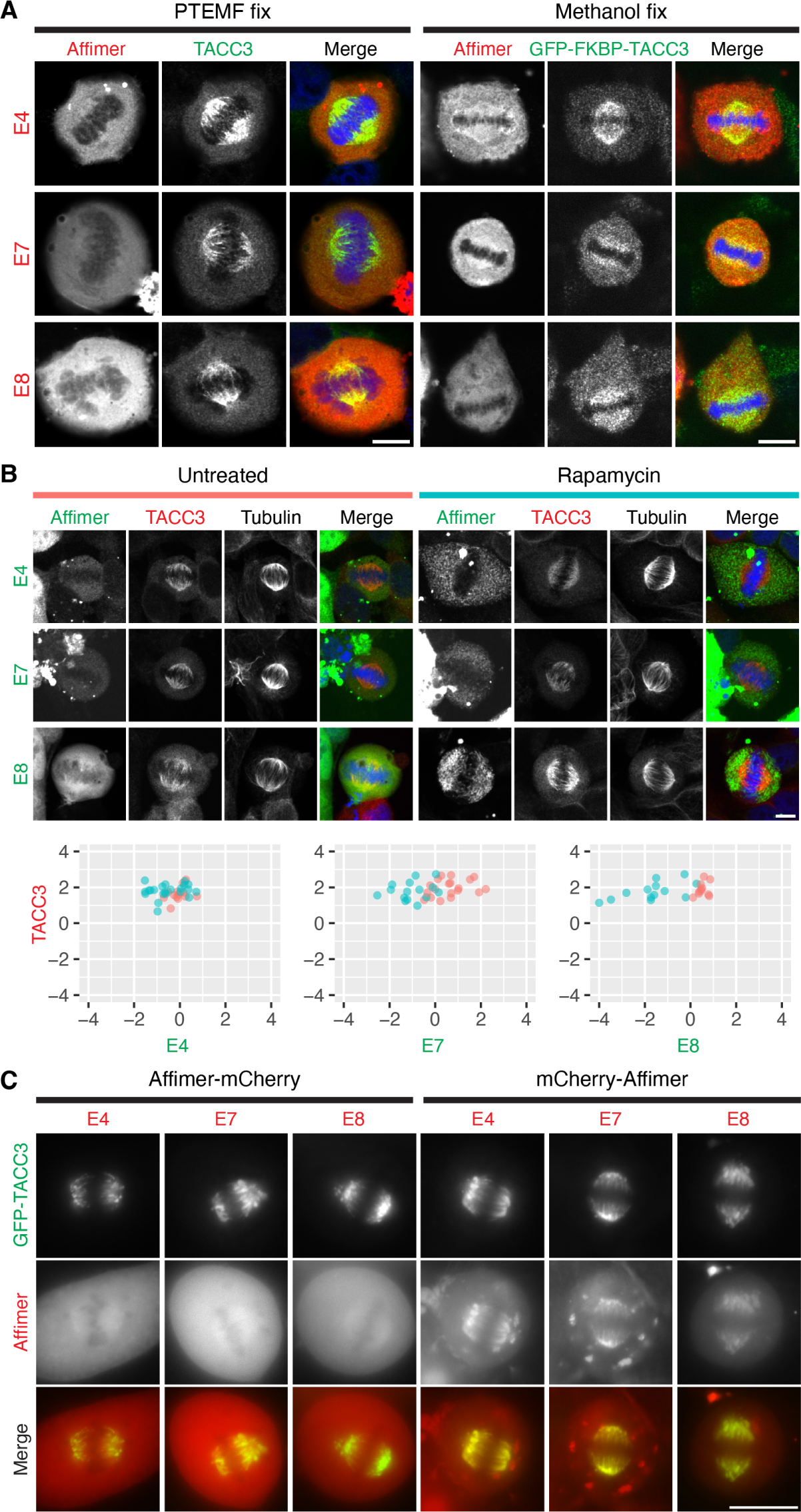
TACC3 Affimers cannot be used to localize or inducibly relocalize endogenous TACC3. (**A**) Representative confocal micrographs of metaphase HeLa cells (left) and CRISPR GFP-FKBP-TACC3 HeLa cells (right) expressing the indicated mCherry-Affimers (red). Cells were fixed with PTEMF or ice-cold methanol, as indicated. Where parental HeLa cells were used, an antibody to stain endogenous TACC3 (green) was used. (**B**) Induced relocalization of TACC3 Affimers to mitochondria. Representative confocal micrographs of HeLa cells at metaphase expressing the indicated FKBP-GFP-Affimers (green) with dark-MitoTrap, that were either treated or not with rapamycin (200 nM, 30 min) prior to fixation. Cells were stained for tubulin (not shown in merge) and TACC3 (red). DNA (blue) is shown in the merge. Relocalization of the Affimer to mitochondria can be seen in the rapamycin-treated cells compared to control, but no relocation of TACC3 is observed. (**C**) Quantification of Affimer (x-axis) and TACC3 (y-axis) spindle localization in untreated cells (salmon) and rapamycin treated cells (turquoise). Spindle localization was calculated as the ratio of spindle to cytoplasmic fluorescence shown on a log2 scale, n = 11-22 cells per condition. (**D**) Widefield micrographs of live HeLa cells in metaphase expressing GFP-TACC3 (green) and TACC3 Affimers (red) labeled with mCherry at either the C or N-terminus as indicated. Scale bars, 10 µm.

**Figure S8.**
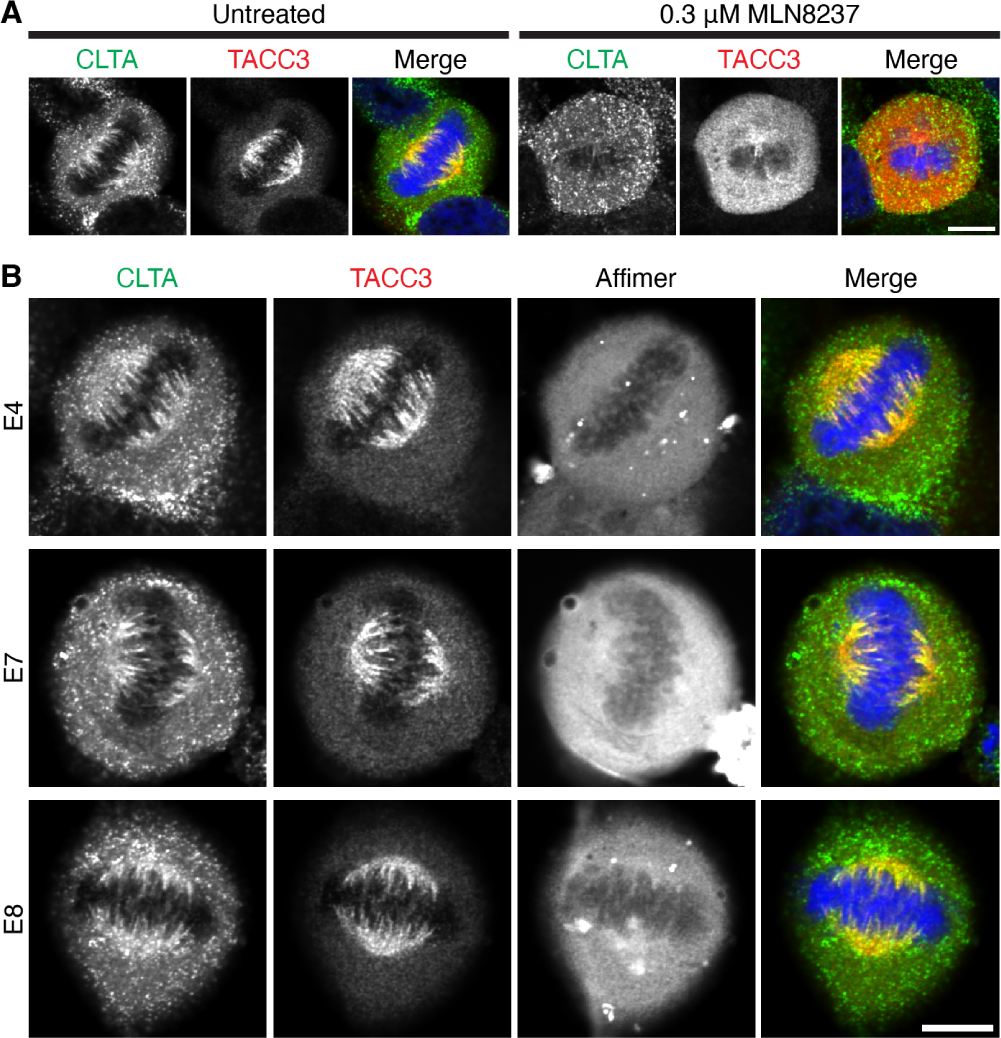
TACC3 Affimers do not affect the mitotic spindle localisation of clathrin or TACC3. (**A**) Representative confocal micrographs of untreated or MLN8237-treated (0.3 µM, 40 min) knock-in CLTA-FKBP-GFP HeLa cells at metaphase. Cells were fixed in PTEMF and stained for TACC3 (red), DNA (blue), and a GFP-boost antibody was used to enhance the signal of CLTA-FKBP-GFP (green). (**B**) Cells expressing the indicated mCherry-Affimers (grey, not shown in merge). Scale bar, 10 µm.

**Figure S9.**
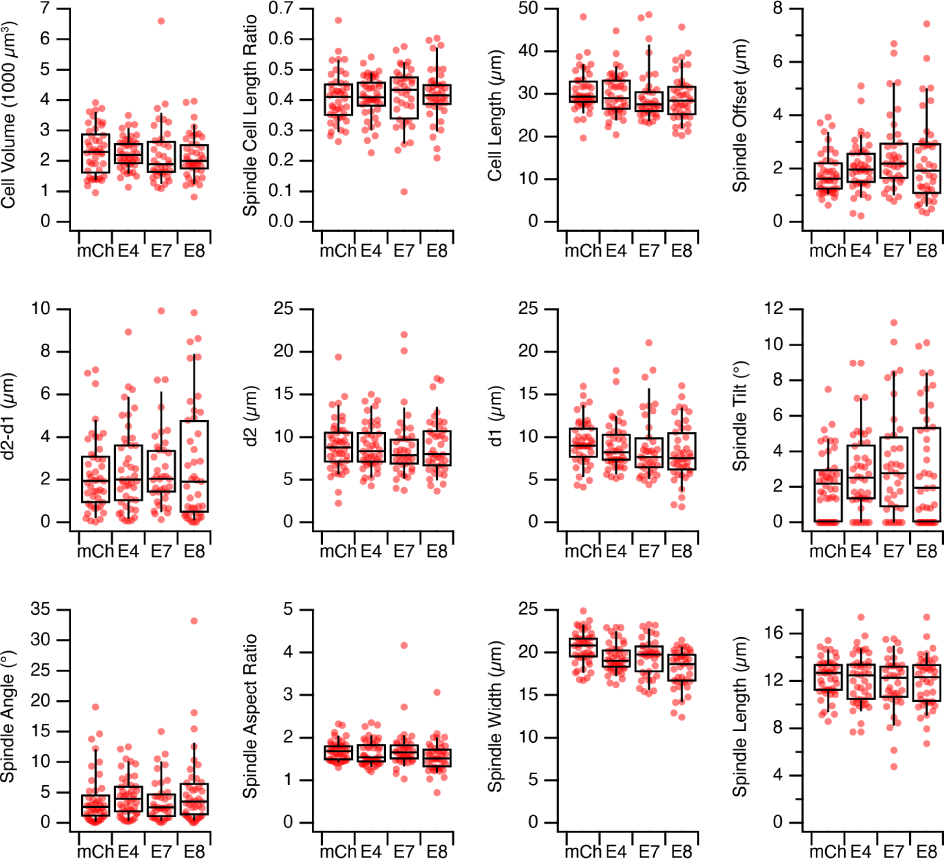
TACC3 Affimers do not affect mitotic spindle morphology or positioning in HeLa cells. Quantification of mitotic spindle parameters in HeLa cells expressing mCherry (mCh) or mCherry-Affimers (E4, E7, E8). Cells were fixed in PTEMF and stained for α-tubulin, pericentrin and DNA. Dot and box plots show the spindle parameters were measured using a semi-automated workflow. Spindle offset is the euclidean distance between the cell center and the spindle center. The distances d2 and d1 refer to the distance from each centrosome to the cell boundary (taken as a sphere that best fit the 3D perimeter of the cell). Spindle tilt and angle are the angle between the spindle axis and the imaging plane or the metaphase plate, respectively. Each dot represents a single cell, n = 37-49 cells per condition pooled from three independent experiments. Box shows the interquartile range, the line the median and the whiskers show the 9th and 91st percentile.

**Figure S10.**
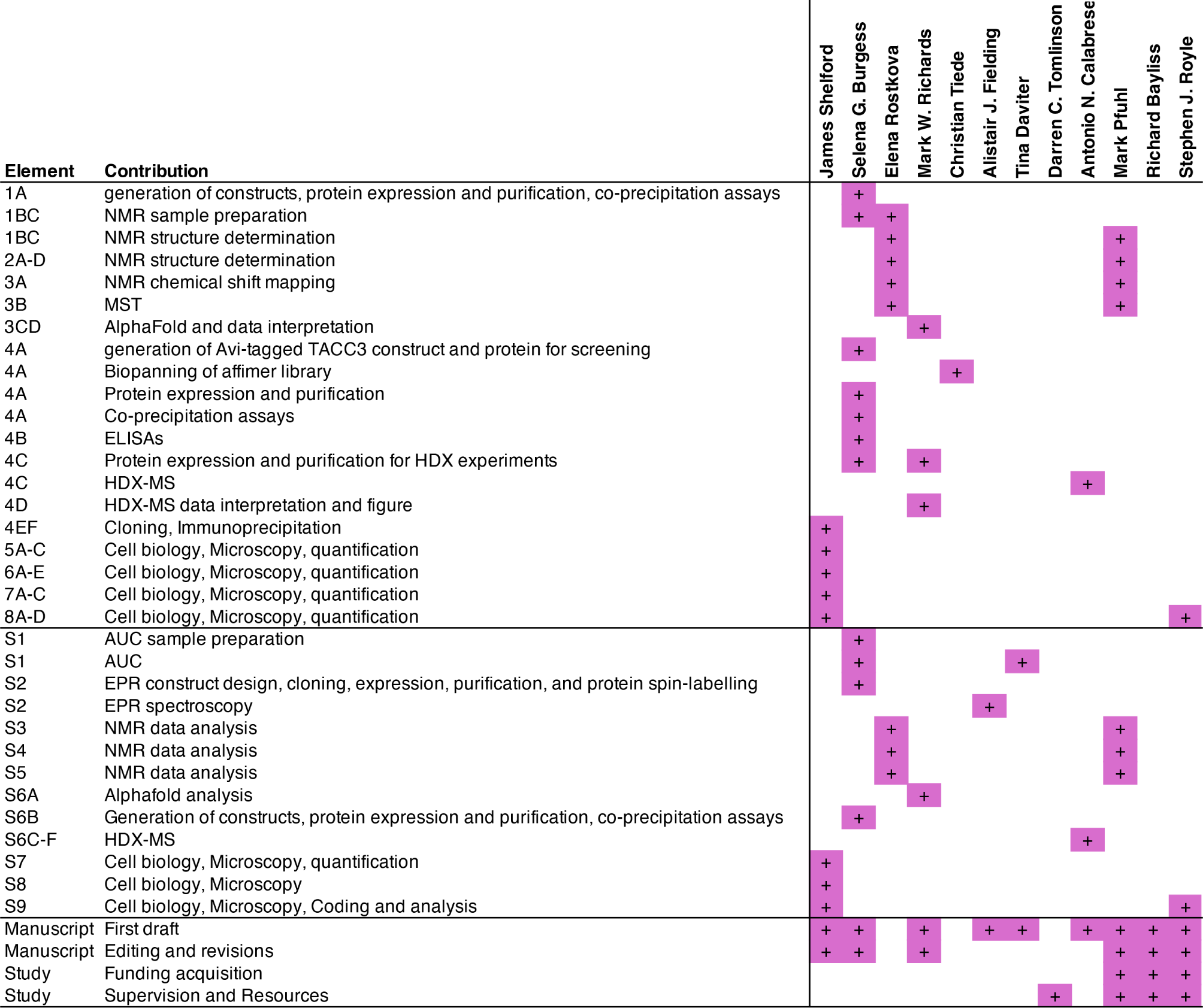
Author contribution matrix.

## Supplementary Tables

**Table S1.**
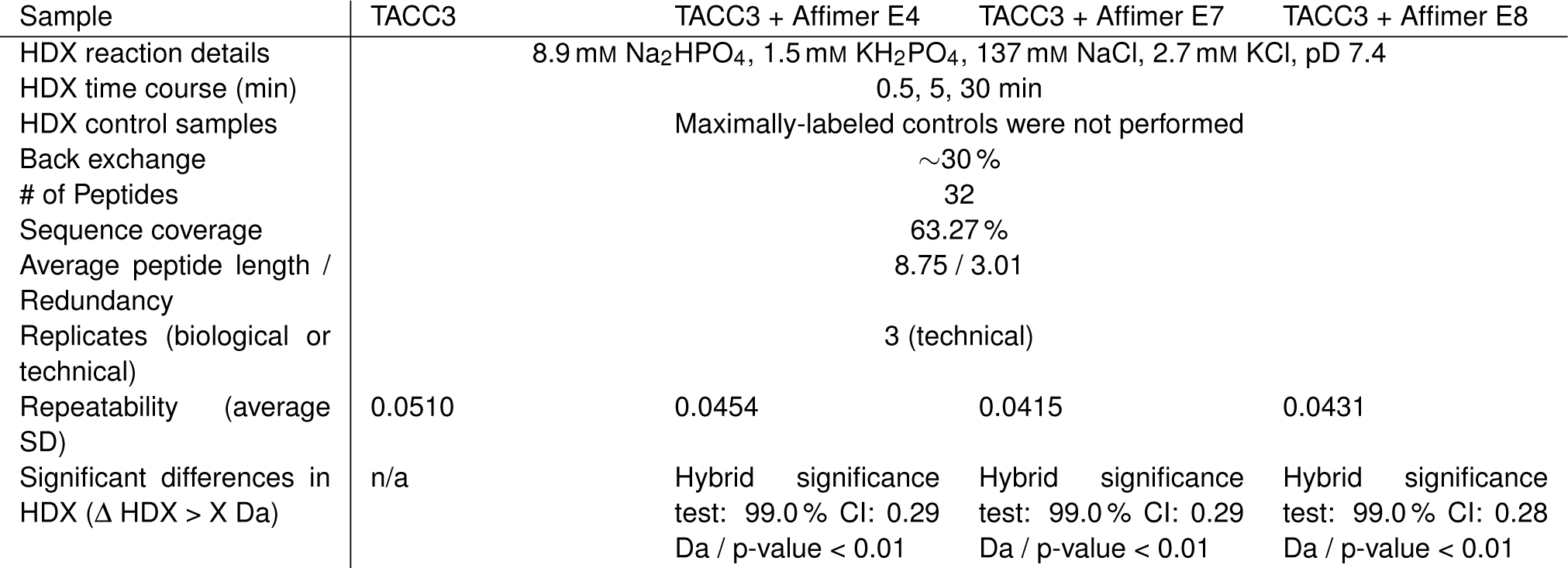
HDX Data Summary Table. SD = standard deviation, CI = confidence interval.

## Notes

https://github.com/quantixed/p062p035

### Summary of Updates

Updated conflict of interest section: "Affimer reagents were invented by DCT and patented by the University of Leeds. The patent was licensed to Avacta Life Sciences, royalties are received as part of this agreement and it is managed by University of Leeds Commercialization Services."

